# Species’ range dynamics affect the evolution of spatial variation in plasticity under environmental change

**DOI:** 10.1101/344895

**Authors:** Max Schmid, Ramon Dallo, Frédéric Guillaume

**Affiliations:** Department of Evolutionary Biology and Environmental Studies, University of Zurich Winterthurerstrasse 190, CH-8057 Zurich, Switzerland.; Masters degree program Biology ETH Zurich, Switzerland

**Author notes:** **Corresponding author:** Fréedéeric Guillaume, Department of Evolutionary Biology and Environmental Studies University of Zurich, Winterthurerstrasse 190, CH-8057 Zurich, Switzerland.

**Keywords:** phenotypic plasticity, environmental tolerance, range shift, environmental change, genetic assimilation, perception trait, trailing edge, leading edge, range expansion

## Abstract

While clines in environmental tolerance and phenotypic plasticity along a single species’ range are widespread and of special interest in the context of adaptation to environmental changes, we know little about their evolution. Recent empirical findings in ectotherms suggest that processes underlying dynamic species’ ranges can give rise to spatial differences in environmental tolerance and phenotypic plasticity within species. We used individual-based simulations to investigate how plasticity and tolerance evolve in the course of three scenarios of species’ range shifts and range expansions on environmental gradients. We found that regions of a species’ range which experienced a longer history or larger extent of environmental change generally exhibited increased plasticity or tolerance. Such regions may be at the trailing edge when a species is tracking its ecological niche in space (e.g., in a climate change scenario) or at the front edge when a species expands into a new habitat (e.g., in an expansion/invasion scenario). Elevated tolerance and plasticity in the distribution center was detected when asymmetric environmental change (e.g., polar amplification) led to a range expansion. Greater gene flow across the range had a dual effect on plasticity and tolerance clines, with an amplifying effect in niche expansion scenarios (allowing for faster colonization into novel environments), but with a dampening effect in range shift scenarios (favoring spatial translocation of adapted genotypes). However, tolerance and plasticity clines were transient and slowly flattened out after range dynamics because of genetic assimilation. In general, our approach allowed us to investigate the evolution of environmental tolerance and phenotypic plasticity under transient evolutionary dynamics in non-equilibrium situations, which contributes to a better understanding of observed patterns and of how species may respond to future environmental changes.

**Impact Summary:** In a variable and changing environment, the ability of a species to cope with a range of selection pressures and a multitude of environmental conditions is critical, both for its’ spatial distribution and its’ long-term persistence. Striking examples of spatial differences in environmental tolerance have been found within species, when single populations differed from each other in their environmental optimum and tolerance breadth, a characteristic that might strongly modify a species’ response to future environmental change. However, we still know little about the evolutionary processes causing these tolerance differences between populations, especially when the differences result from transient evolutionary dynamics in non-equilibrium situations. We demonstrate with individual-based simulations, how spatial differences in environmental tolerance and phenotypic plasticity evolved across a species’ range during three scenarios of range shifts and range expansion. Range dynamics were either driven by environmental change or by the expansion of the ecological niche. The outcome strongly differed between scenarios as tolerance and plasticity were maximized either at the leading edge, at the trailing edge, or in the middle of the species’ range. Spatial tolerance variation resulted from colonization chronologies and histories of environmental change that varied along the range. Subsequent to the range dynamics, the tolerance and plasticity clines slowly leveled out again as result of genetic assimilation such that the described responses are long-lasting, but in the end temporary. These findings help us better understand species’ evolutionary responses during range shifts and range expansion, especially when facing environmental change.

## Introduction

Species exhibit remarkable abilities to survive in variable environments, which manifests as environmental tolerance, and has caught the interest of biologists early on (e.g., Grinnell, 1917; Elton, 1927; Hutchinson, 1957). A species’ environmental tolerance can be broadly defined as its ecological niche, an important conceptual tool to understand the geographical distribution of species (Guisan and Zimmermann, 2000; Essl et al., 2009; Slatyer et al., 2013) and to predict their response to environmental changes (Hijmans and Graham, 2006; Valladares et al., 2014). At a lower level, environmental tolerance can be related to the capacity of a genotype to produce a plastic phenotype and to adapt to varying environmental and ecological conditions (Chevin et al., 2010). The capacity of a genotype to produce adapted phenotypes is, however, rarely perfect and organisms often have an environment in which they perform best (e.g., Eppley, 1972; Huey and Kingsolver, 1989). The relationship between environmental variation and organismal performance thus often results in a bell-shaped curve with performance decreasing away from an optimal environmental condition. Yet, not only do species differ in the breadth of their niche, they also differ in their environmental tolerance and plasticity among populations within their range (e.g., Macdonald and Chinnappa, 1989; Woods et al., 2012; Bennett et al., 2015; Lancaster, 2016; Toftegaard et al., 2016; Reger et al., 2018). For example, terrestrial ectotherms often exhibit clines in environmental tolerance (including arthropods, amphibians, and reptiles) with broader thermal tolerances at high latitudes compared to species or populations near the equator (Addo-Bediako et al., 2000; Gaston, 2009; Sunday et al., 2012; Lancaster et al., 2015; Lancaster, 2016). While inter- and intra-species spatial variation in tolerance breadth and plasticity are well documented, a comprehensive understanding of the underlying biological processes has not been achieved yet.

Divergence in tolerance and niche breadth can result from evolutionary processes, given that environmental tolerance is a heritable trait with additive genetic variance for fitness (Via and Lande, 1985; Lynch and Gabriel, 1987; Chevin and Lande, 2011). Selection pressures affecting tolerance evolution mainly stem from the environmental variability that genotypes experience, across space or over time. In temporally variable environments (Lynch and Gabriel, 1987; Lande, 2014), or in structured populations with gene flow between distinct habitats (Via and Lande, 1985; Sultan and Spencer, 2002; Van Buskirk, 2017), environmental tolerance is expected to evolve as a means to increase the average fitness of genotypes encountering multiple habitats. In contrast, low levels of environmental variability lessen the fitness benefits of a broad environmental tolerance and limit its evolution when a trade-off is present (Lynch and Gabriel, 1987; Padilla and Adolph, 1996; Reed et al., 2010; Ezard et al., 2014). Maintaining environmental tolerance via phenotypic plasticity may also entail physiological or metabolic costs, further opposing its evolution (Moran, 1992; van Buskirk and Steiner, 2009; Lande, 2014). When the environmental variability differs between populations within a species, we can thus expect spatial differences in tolerance to evolve. For example, following the observation that temperature vary more strongly across a day or a season at high than low latitudes, the *climate variability hypothesis* attempts to explain latitudinal differences in ectotherms’ thermal tolerance as a consequence of the observed spatial difference in temperature fluctuations (Janzen, 1967; Stevens et al., 1989; Gaston and Chown, 1999), with higher thermal tolerance selected in northern habitats. However, while some empirical studies seem to support the climate variability hypothesis (Addo-Bediako et al., 2000; Vázquez and Stevens, 2004), others suggest that alternative mechanisms could cause the same spatial differentiation of environmental tolerance, especially range expansion (Lancaster, 2016).

Range expansion, as when species invade new habitats, or during post-glacial migration, can cause a temporary and local increase in environmental tolerance creating clinal patterns of plasticity and tolerance across species’ ranges. For example, in a recent meta-analysis, Lancaster (2016) showed that higher thermal tolerances at high latitudinal margins were only found for insect species that are currently or were recently expanding or shifting their range towards the poles. Instead, ectotherms with stable distributions, mostly endemic or insular species, were shown to have constant thermal tolerance breadths across latitudes. Lancaster (2016) concluded that temporary evolutionary dynamics in the course of range shifts or range expansions are responsible for observed latitudinal clines in thermal tolerance breadth. Such temporary dynamics have been found in analytical models where a transient increase of adaptive phenotypic plasticity appears in populations facing temporal changes of their local environment and thus increase their environmental tolerance to better cope with novel environments (Gavrilets and Scheiner, 1993; Lande, 2009; Chevin and Lande, 2011; Gallet et al., 2014). Following stabilization of the environment, genetic assimilation can cause a reduction of tolerance and phenotypic plasticity, and lead to canalization of the genotypes (Waddington, 1953; Crispo, 2007; Lande, 2009; Ergon and Ergon, 2016). Similarly, expanding a species’ range by adapting to novel environmental conditions outside of the current ecological niche can be achieved by a transient increase in plasticity and tolerance in the newly founded populations (Lande, 2009; Chevin and Lande, 2011; Lande, 2015). Plasticity clines can thus have two different origins: range dynamics or clines of climate variability. No attempt has yet been made to distinguish between these two causes, and to investigate the effect of range dynamics on tolerance and plasticity evolution in detail.

Species’ range dynamics can result from the colonization of novel habitats and the evolution of a species’ ecological niche. Niche expansion can be achieved when a species evolves either its environmental optimum, its tolerance breadth or both in novel habitats (Wilson, 1961; Thomas et al., 2001; Wiens and Donoghue, 2004; Early and Sax, 2014; Atwater et al., 2017). Niche evolution is an important driver of invasive species’ range dynamics when the evolution of increased tolerance allows alien species to become invasive in the novel habitat (Brock et al., 2005; Richards et al., 2006; Lande, 2009; Alexander and Edwards, 2010; Chevin and Lande, 2011; Lande, 2015). However, the evolution of the ecological niche is not a necessary prerequisite for changing species’ ranges. Large-scale environmental changes can force a species to track its suitable environmental conditions in space and shift its range accordingly. This scenario of range shift may be valid for latitudinal shifts of species’ ranges after the last ice age (Hewitt, 1999, 2000), or for more recent responses to global climate warming (Parmesan, 2006; Tingley et al., 2009; Talluto et al., 2017). Alternatively, a range expansion may occur when the rate of environmental change is not constant across space, as when global temperature changes much faster at high than low latitudes (known as polar amplification, see box 5.1 in Stocker et al., 2013). Populations at the northern range margin may thus follow rapidly shifting local conditions and expand into new geographical areas while trailing edge populations at the southern range margin may face slower local changes and rather stay in place, while keeping the 6 species’ niche constant. Spatial plasticity clines may here differ from scenarios with uniform rates of environmental change or from scenarios of niche evolution in a constant environment. In short, changes in the species’ distribution can be triggered by the evolution of the species’ ecological niche and by environmental change. While plasticity and tolerance evolution as drivers of range expansions have been studied before in two-patch models (Lande, 2015), theoretical work is missing that approaches plasticity and tolerance evolution during large-scale environmental change and comprises an entire species’ distribution. Here, a better understanding might not only allow to unravel past processes underlying large-scale biogeographic patterns, but also better predict future evolutionary responses to environmental change.

In this paper, we make a distinction between range shift, range expansion, and niche expansion and show that these three evolutionary scenarios differ in the resulting spatial 8 clines of environmental tolerance and phenotypic plasticity. We used two different approaches of modeling environmental tolerance, working with an evolving tolerance curve and evolving norm of reactions. Using individual-based simulations, we show that varying 1 histories of environmental change in local populations set on an environmental gradient can act as a driver of tolerance differentiation between populations, even in absence of a species’ niche expansion and spatial differences in environmental variability. We further found that plasticity clines can be in opposite directions depending on whether a species expands its niche into new habitats or follows it across space.

## Methods

To model the evolution of environmental tolerance, we used two common approaches, a tolerance curve and a norm of reaction. The tolerance curve describes fitness in dependence of the environment in a very general way (Lynch and Gabriel, 1987), without an environment dependent phenotypic representation (Lande, 2014). Alternatively, modeling phenotypic plasticity using a genotypic reaction norm explicitly maps phenotypes to environments (Via et al., 1995; Whitlock, 1996; Chevin et al., 2010; Svanbäck and Schluter, 2012; Lande, 2014; Valladares et al., 2014). We used individual-based simulations with a modified version of Nemo (Guillaume and Rougemont, 2006) to model evolving tolerance curves and reaction norms.

### Tolerance curve

We modeled the tolerance curve as a Gaussian function describing fitness (i.e., survival probability) of a phenotype in dependence of the environment (see Fig. 1a). The tolerance optimum (*t*_0_, the phenotype) and tolerance breadth (*t*_1_) were determined by two evolving quantitative traits. We implemented a generalist-specialist trade-off by imposing a constraint between the height and the breadth of the tolerance curve where the maximum fitness *W_max_*(*t*_1_) is a decreasing function of the tolerance breadth (*t*_1_) (see Fig. 1a and equation 5). The fitness of an individual with tolerance traits *t*_0_ and *t*_1_ has fitness *W*(*e*) when responding to the environment *e*:
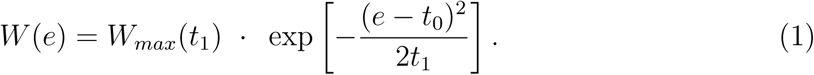

**Figure 1:**
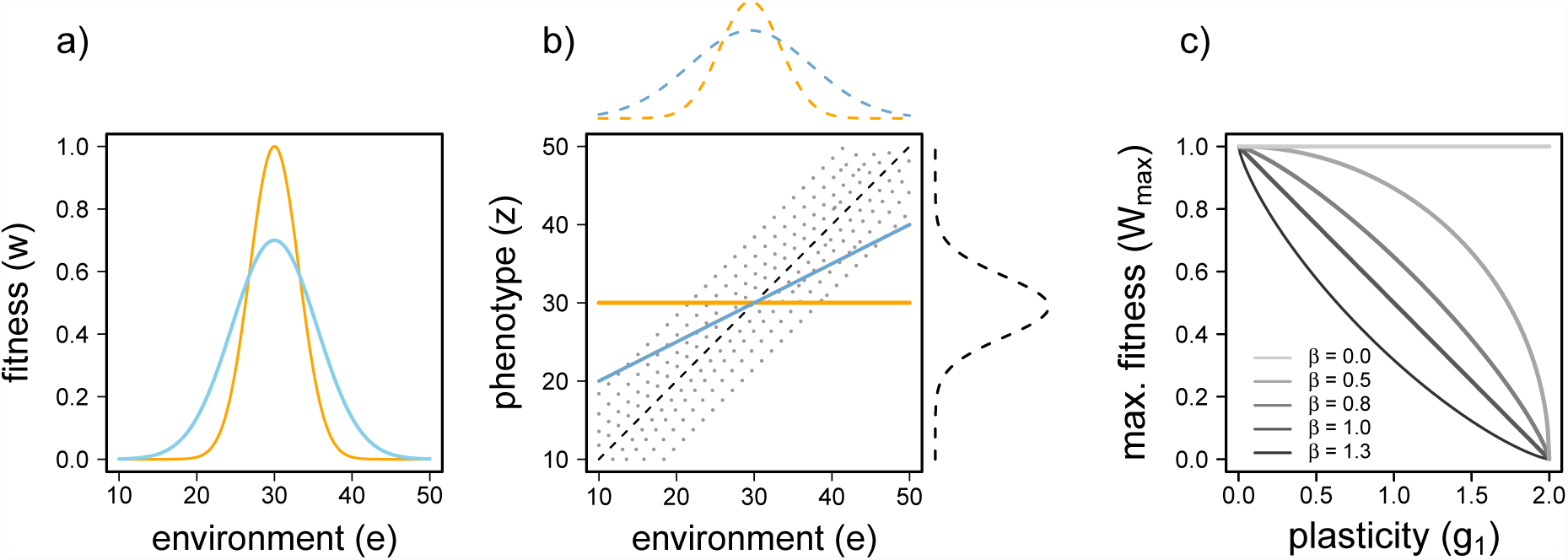
Two tolerance curves are illustrated in graph a) with identical environmental optima (*t*_0_ = 30), but different tolerance breadths (*t*_1_). Given a specialist-generalist trade-off (*β* > 0, see Equation 4), a higher tolerance translates into a lower maximal fitness. Graph b) illustrates how phenotypic plasticity translates into environmental tolerance. The two solid lines in blue and orange represent two genotypes as linear reaction norms that describe phenotype expression in dependence of the environment. The two genotypes exhibit different degrees of plasticity (orange - no plasticity; blue - adaptive plasticity). The dotted lines in gray show the fitness landscape with maximum fitness achieved at the black dashed line representing the position of the phenotypic optimum O (here assuming Θ = *e*). The two reaction norms translate into two different tolerance curves depending on the amount of plasticity (colored dashed lines at the top of the graph). The black dashed line to the right represents the fitness function (i.e., fitness in dependence of the phenotype) at e = 30. Graph c) shows the costs of plasticity (*β*) that reduces the maximal fitness (*W_max_*) when the absolute value of plasticity deviates from zero.

### Phenotypic plasticity

We implemented phenotypic plasticity as a linear norm of reaction (NoR; see Fig. 1b), as in Schmid and Guillaume (2017) (see also Scheiner, 1998; Scheiner et al., 2012). The phenotype of each individual (*z*) is expressed in dependence of its genotype and the environment in which it develops. The genotype codes for two evolving quantitative traits, the NoR intercept (*g*_0_) and the NoR slope (*g*_1_). The environment then affects the phenotype depending on the environmental deviation from the reference environment (*g*_2_), also called the perception trait (Lande, 2009; Ergon and Ergon, 2016). We kept the perception trait value constant. The environment-dependent trait value *z*(*e*) is then:
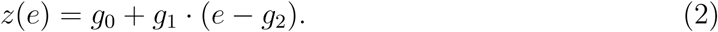

The NoR intercept *g*_0_ relates to the genotypic value measured in the reference environment *e* = *g*_2_, where the effect of plasticity cancels out (i.e., *g*_1_ · (*e* − *g*_2_) = 0) and only *g*_0_ contributes to the phenotype. The NoR slope *g*_1_ controls the degree of plasticity, that is, how strongly the expressed phenotypes differ between environments. Because the environmental position of the perception trait is a key factor in reaction norm evolution (Ergon and Ergon, 2016, see also Appendix 1), we ran simulations with three different values of *g*_2_ (*g*_2_ = 10, 20, 30).

After the phenotype has been expressed based on the reaction norm, we used a Gaussian selection function to determine the absolute fitness value *W*(*z*) as the individuals’ survival probability (Fig. 1b, black dashed line). *W*(*z*) is a Gaussian function of the distance between the expressed phenotype (*z*(*e*)) and the phenotypic optimum (Θ), depending on the strength of selection (inversely related to the width of the Gaussian curve *ω*^2^) and the cost of plasticity (*W_max_*(*g*_1_)):
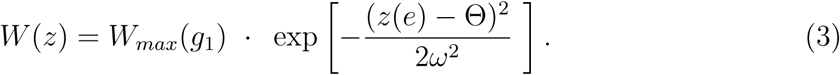

### Costs of plasticity and the generalist-specialist trade-off

We modeled constitutive costs of plasticity (sensu Chevin et al., 2010) such that *W_max_* declines with increasing (absolute) values of the NoR slope (*g*_1_) (Fig. 1c), or tolerance breadth (*t*_1_). For the cost of plasticity in the NoR model, we used a modified version of the trade-off function from Débarre and Gandon (2010):
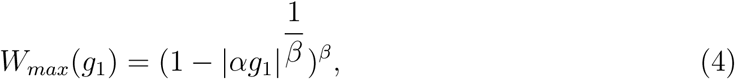

with the scale parameter *α*, and the shape parameter *β* which controls the concavity of the relationship. While *α* was 0.5 in all of our scenarios, we ran simulations with 5 different values of *β* (*β* = 0.0, 0.5, 0.8,1.0,1.3). In absence of costs (*β* = 0.0), maximum fitness is high for all values of plasticity, while *β* > 0 causes a negative relationship between plasticity and maximal fitness, installing a specialist-generalist trade-off. The higher *β* the more constrained the evolution of plasticity (Fig. 1c). Furthermore, we explored the effect of three different selection strength values (*ω*^2^ = 1, 4,16).

For a better comparison with the evolution of phenotypic plasticity, we translated the cost of plasticity linked to *g*_1_ into the cost of tolerance represented by its breadth parameter 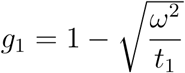 (for the derivation see Appendix 2) such that:
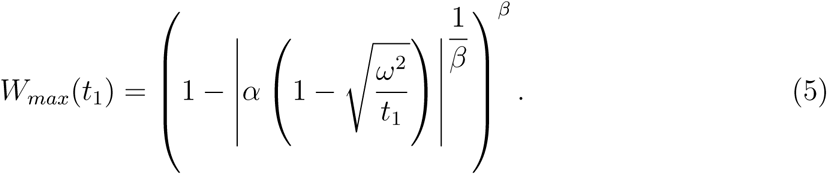

### Scenarios of species’ range evolution

#### Initial environmental conditions (burn-in simulations)

We modeled 42 habitat patches that were linearly arranged along an environmental gradient connected by nearest-neighbor dispersal (i.e., stepping stone model). Average environmental values per patch (*e*) ranged from 10 at the left margin to 51 at the right margin with a constant between-patch environmental distance of 1. The phenotypic optima were identical to the environmental values (Θ = *e*), we will thus only refer to e from now on. In the burn-in, we set the initial range within the first 21 patches on the gradient, with environmental values between 10 ≤ *e* ≤ 30, and a carrying capacity of 200 individuals (see Fig. 2a, dashed gray line). Patches to the right of the initial range were designated as not habitable and attributed a carrying capacity of zero. Environmental conditions also varied randomly within patches across generations and individuals such that the environmental value experienced by each individual (and thus the phenotypic optimum Θ) was picked from a Gaussian distribution with mean e and variance *σ*^2^(*e*) = 1. Consequently, a population experienced within-patch environmental variation either resulting from spatial heterogeneity within a patch or from temporal fluctuations when individuals were born and experienced selection at slightly different points in time. For each parameter combination (Tab. 1) we ran burn-in simulations for 100,000 generations and 20 replicates on a constant average environmental gradient to reach migration-selection-drift balance.

**Figure 2:**
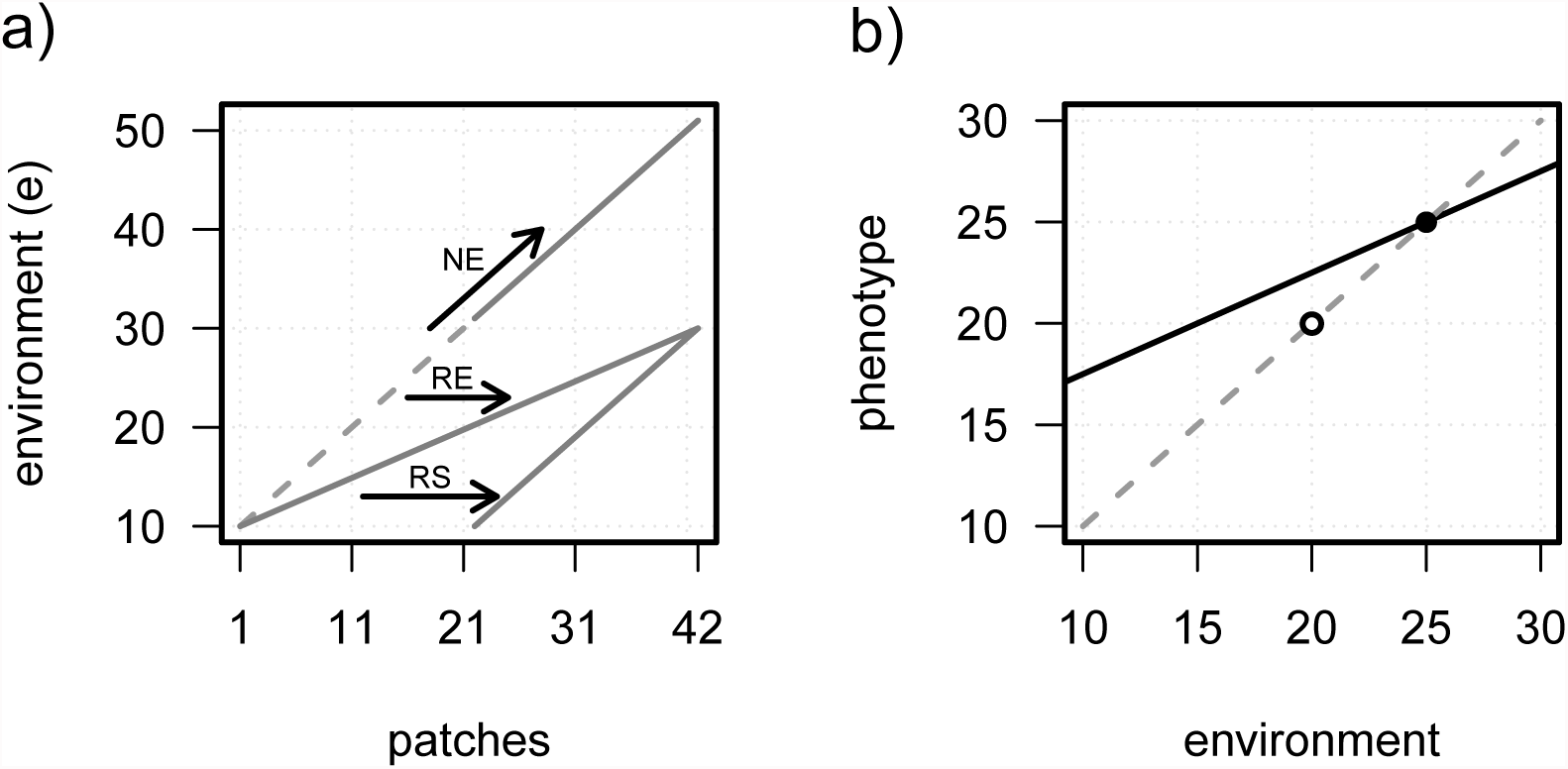
Cartoons showing the three scenarios of species’ range shifts and environmental change. In a), 42 patches (*x*-axis) are linearly arranged along an environmental gradient (*y*-axis) with environmental values *e* (= phenotypic optima Θ). The dashed line shows the environmental values set in the burn-in simulations before range expansion (21 patches on the left). The solid lines show the environmental values in patches at the end of the expansion/shift simulations after 210 generations for each scenario (RS: range shift, RE: range expansion, NE: niche expansion, see text for details). Within a single patch, b) illustrates the average NoR (black line), the phenotypic optima (gray dashed line) and the experienced environment before (solid dot) and after (circle) environmental change in RS and RE scenarios. The lower phenotypic optimum is partially reached thanks to an adaptive plastic response (positive NoR slope).

**Table 1:** The following table shows the explored parameter space of the burn-in and range dynamic simulations (RS, RE, NE). We run 20 replicates for every parameter combination.

| Parameter | Values |
| --- | --- |
| costs of plasticity ($\beta$) | 0.0 , 0.5 , 0.8, 1.0 , 1.3 |
| migration rate ($m_c$) | 0.001, 0.01 , 0.1, 0.2 , 0.4 |
| selection strength ($\omega^2$) | 1 , 4 , 16 |
| perception trait value ($g_2$) | 10 , 20 , 30 |

#### Range shift (RS)

After burn-in, the average environmental conditions within patches started to change with rate Δ*e* = −0.1 per generation (Fig. 2a). As we did not allow the ecological niche to evolve in this scenario, patches with environmental values outside the initial species’ range were not available for colonization and their carrying capacities were set to zero. Therefore, patches at the trailing edge became inhabitable when their local environmental value dropped below 9.5. Alternatively, new patches at the front edge became available for colonization when their environmental value reached 30.5. With rates Δ*e* = −0.1, the species’ niche shifted completely to the right of the environmental gradients and settled into patches 22–42 in 210 generations.

#### Range expansion (RE)

In this scenario, we explored the consequences of range expansion while maintaining the ecological niche constant. We achieved this by setting variable rates of environmental change across the range, starting with a constant habitable environment at the left margin (constant edge) and a maximum rate of change at the expanding edge of the range (set at Δ*e* = −0.1), until patch 42 reached an environmental value of 30.5 and became habitable (Fig. 2a). The rate of change in the rest of the range was linearly decreased to maintain a linear environmental gradient among patches as the range increased. This scenario mimics environmental change with an extreme case of polar amplification (Stocker et al., 2013, Box 5.1), when environmental change (e.g., global warming) is stronger at one edge of the gradient (at high latitudes or altitudes) compared to the other edge (at the equator and low altitudes). In this model, new patches became habitable every 10 generations at the right margin. The time course was similar to RS with Δ*e* = −0.1 such that patch 42 became habitable after 210 generations of range expansion.

#### Niche expansion (NE)

Finally, we modeled niche expansion on a constant environmental gradient by allowing for range expansion into novel habitats. After the burn in, the individuals were allowed to colonize the 21 new habitat patches on the right (Fig. 2a). To allow for a better comparison with RS and RE scenarios, the patches on the right were opened for colonization in a stepwise fashion every 10 generations, with the last patch becoming habitable after 210 generations, insuring that the total time for colonization was the same as with Δ*e* = −0.1.

In all three scenarios, we have run the simulations for an additional 180 generations with stable environmental conditions to study genetic assimilation.

### Genetic parameters

Individuals were diploid in random mating populations. The two evolving quantitative traits were coded on 20 unlinked quantitative trait loci (QTL), each allele contributing additively and pleiotropically to two traits (tolerance curve optimum *t*_0_ and breadth *t*_1_, or NoR intercept *g*_0_ and NoR slope *g*_1_). We used a mutation rate of *m* = 0.0001 per allele and a continuum-of-allele mutation model where mutational effects were picked from an uncorrelated bivariate normal distribution centered on (0, 0) and added to the standing allelic effects. Mutational variance *α*^2^ was set to 0.1 for (*g*_0_) and 0.001 for (*g*_1_) in the NoR simulations, and to 0.1 for (*t*_0_) and 1 for (*t*_1_) for tolerance curve simulations. We set the mutational covariance to zero (see Fig. A1). We used a higher mutational variance for *t*_1_ than for *g*_1_ as mutational phenotypic effects in *g*_1_ are environment-dependent and increase with the distance between *e* and *g*_2_ along the gradient (see Equation 2, and Fig. A1).

### Life cycle

In all simulations, dioecious individuals mated within patches at random, and without selfing. The female fecundity was picked from a Poisson distribution with a mean of 5, independent of its fitness. Generations were non-overlapping with even sex ratio on average. Each generation started with *breeding* (when adults produced offspring), followed by *migration* (of the offspring), *phenotype expression* and *fitness determination* (of offspring in dependence of the respective environment), *selection* (removing individuals in dependence of their fitness), *regulation* (all adults died; offspring were discarded randomly until carrying capacity was reached) and *aging* (offspring was transferred into adult life stage). As result of this life cycle, plasticity corresponded to developmental or one-shot plasticity (Lande, 2015) when phenotypes were expressed once after migration (e.g., seed dispersal) based on the environment of selection with a perfect reliability of the environmental cue. We modeled the connectivity between patches as a stepping stone model with dispersal only between neighboring patches and absorbing boundaries at the range margins. Simulations were run with six different migration probabilities (*m_c_* = 0.001, 0.01, 0.1, 0.2, 0.4).

## Results

### Static range (burn-in)

After 100,000 generations in a constant environment, average environmental tolerance (*t*_1_) and phenotypic plasticity (*g*_1_) evolved to uniform values along the species’ range, for most parameter combinations (e.g., see Fig. 3 – 5, dashed lines). Lower levels of plasticity and tolerance were observed at the range edges compared to the center when migration was high, because genotypes in the edge populations experienced lower environmental variation across generations due to the absorbing boundaries (Fig. S1).

**Figure 3:**
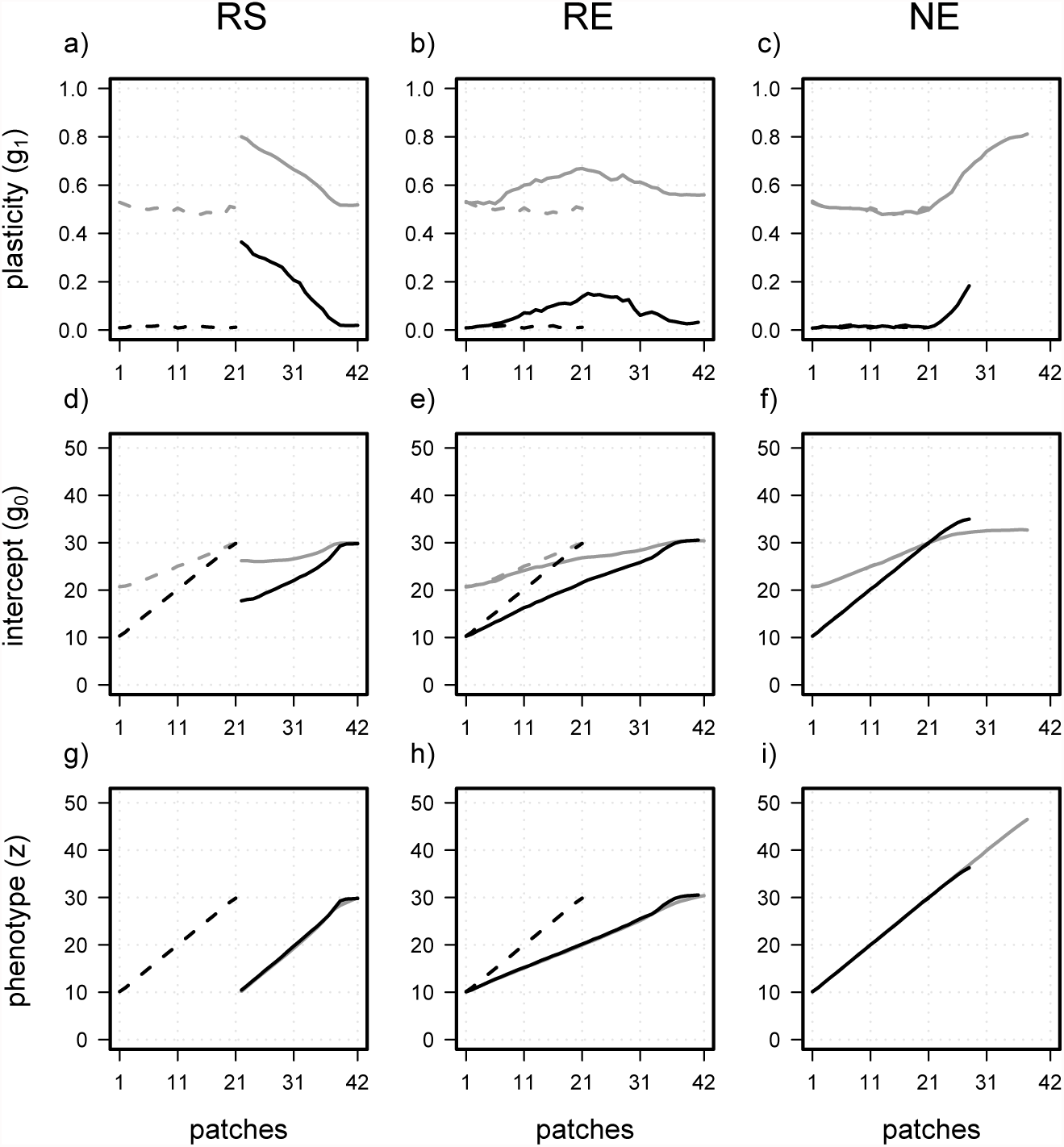
The NoR intercept, NoR slope and the resulting phenotype are shown for scenarios with *ω*^2^ = 4, *σ*^2^ (e) = 1, *m_c_* = 0.01, *g*_2_ = 30 for two different costs (*β* = 0.5 - gray; *β* = 1.0 - black) before (dashed lines) and after (solid lines) environmental change (RS, RE; Δ*e* = −0.1) or niche evolution (NE).

**Figure 4:**
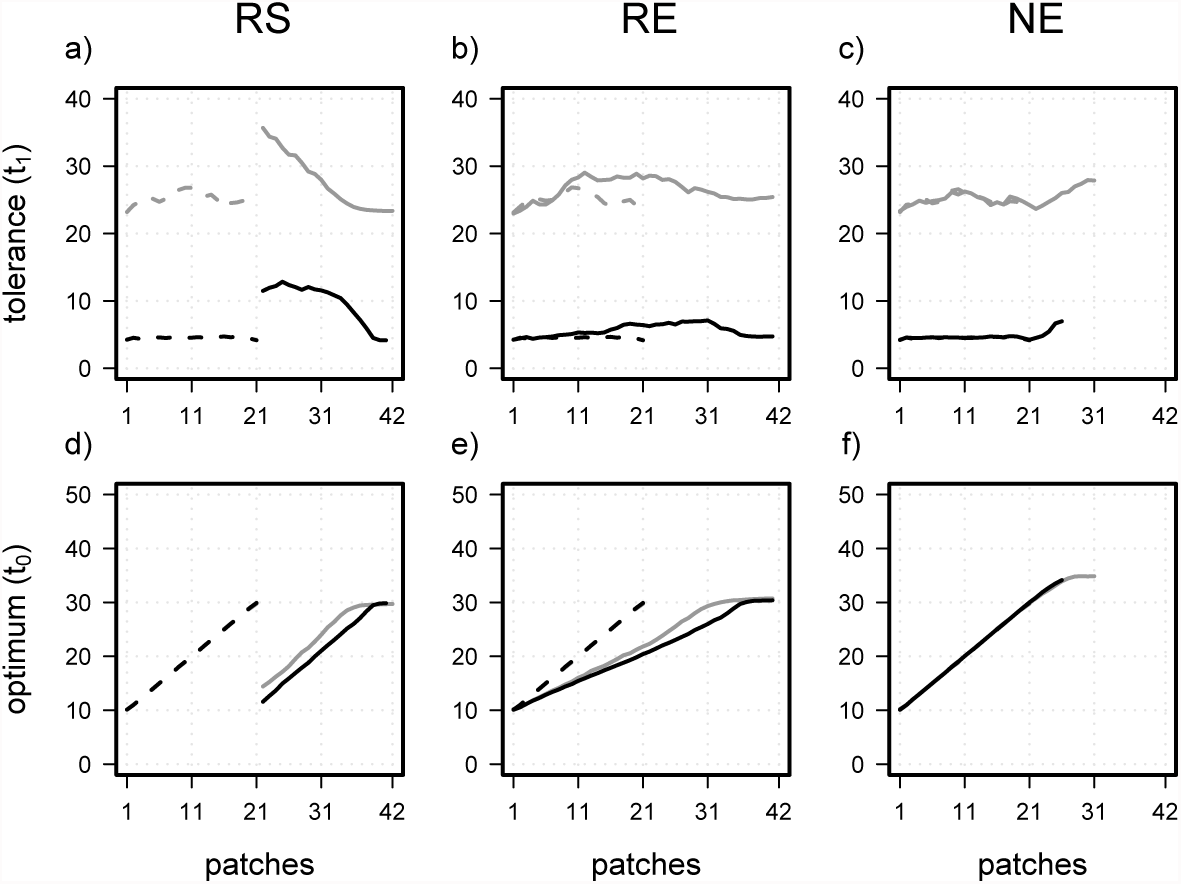
The tolerance breadth and the tolerance optimum for scenarios with *ω*^2^ = 4 and *σ*^2^ (*e*) = 1 for two different costs (*β* = 0.5 - gray; *β* = 1.0 - black) before (dashed lines) and after (solid lines) environmental change (RS, RE; Δ*e* = −0.1) or niche evolution (NE).

**Figure 5:**
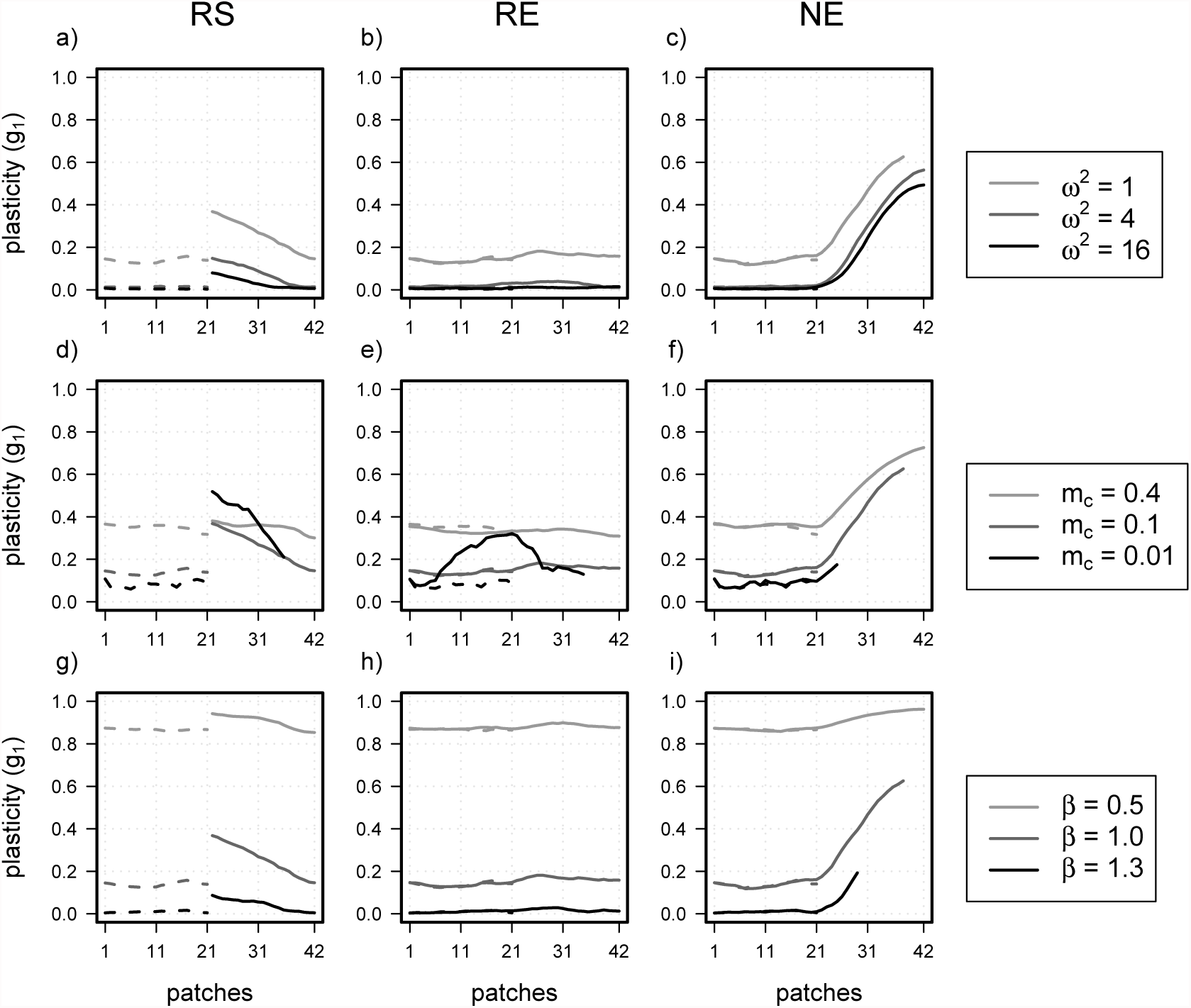
Effects of the strength of selection (*ω*^2^; a-c), dispersal rate (*m_c_*; d-f) and costs (*β*; g-i) on the plasticity clines (*g*_1_). Average phenotypic plasticity per patch is given before (dashed lines) and after (solid lines) the range shift scenarios after 210 generations. Unless specified, results are given for high costs (*β* = 1.0), intermediate environmental variance (*σ*^2^(*e*) = 1), moderate migration rate (*m_c_* = 0.1) with a perception trait value of *g*_2_ = 30, and strong selection (*ω*^2^ = 1).

The average plasticity and tolerance increased with higher migration rate (*m_c_*), stronger selection (low *ω*^2^ values), and lower costs (low *β* values) (Fig. 5, Tab. S1). In absence of costs (*β* = 0), plasticity evolved to be”perfect” (*g*_1_ = 1) and tolerance reached high values (*t*_1_ > 100) (Fig. S2). Lowest values of plasticity and environmental tolerance in absence of costs were achieved with lowest migration rates. The perception trait value (*g*_2_) had no effect on the evolved level of plasticity at equilibrium (Fig. S3).

As expected with increasing levels of average plasticity, the genotypic clines in *g*_0_ across the range were shallower than the phenotypic clines, which matched the environmental values (Fig. 3, Fig. S4a). Phenotypic clines across the range were steeper with high phenotypic plasticity (Fig. S5). A positive covariance between NoR slope (*g*_1_) and NoR intercept (*g*_0_) within populations evolved in all three scenarios after 100’000 generations of burn-in, with covariances decreasing towards the range edges (Fig. S6).

### Environmental change

We observed the evolution of a spatial cline in tolerance and plasticity in all three scenarios of dynamic species’ ranges, although the cline orientation strongly differed between scenarios. In the range shift scenarios (RS), a negative cline evolved with highest plasticity and tolerance at the *trailing* edge of the distribution (Fig. 3a, Fig. 4a, Fig. S7a,d). In range expansion scenarios (RE), plasticity and tolerance were maximized in the middle of the distribution range (Fig. 3b, Fig. 4b, Fig. S7b,e), while in the niche expansion scenarios (NE), a positive cline evolved with the highest values at the expansion front (Fig. 3c, Fig. 4c, Fig. S7c,f).

Environmental tolerance and phenotypic plasticity simulations resulted in qualitatively similar patterns, while slight quantitative deviations were observed, when the evolved plasticity clines were steeper than the tolerance clines (compare Fig. 3a-c to Fig. 4a-c). The evolution of steeper clines in plasticity resulted from mutational effects and additive genetic variance that were environment-dependent for *g*_1_ (and thus varied along the species’ range) but not for *t*_1_. Plasticity thus evolved more than tolerance (*t*_1_) in environments very different from *g*_2_, despite the higher mutational variance in *t*_1_.

In line with expectation of genetic assimilation, the gradient in average tolerance and plasticity leveled out again after range shifts and range expansions (Fig. S8), a process that is 1 slower than the rise of spatial differences in *t*_1_ and *g*_1_ during the range shift (Lande, 2009).

The clines in *g*_1_ and *t*_1_ along the species’ range were steepest, when (1) the strength of selection was high (low *ω*^2^; e.g., Fig. 5a-c), (2) costs were high and initial *g*_1_ and *t*_1_ were small (Fig. 5g-i; Table S2), and (3) migration rate was low (Fig. 5d,e). Lowering the migration rate also strongly reduced the rate of range expansion, especially in the NE scenario (Fig. 5f). No clines in plasticity or tolerance evolved in the RE scenario except for small migration rates (*m* ≤ 0.01). Steeper plasticity clines under low migration rates can be explained by a reduced migration load on plasticity evolution and a reduced translocation of genotypes from neighboring populations with adaptive *g*_0_ and *t*_0_ values.

Plasticity clines were shallower in RE than in RS scenarios because of the smaller extent of environmental change across the range and within patches in RE relative to RS (see Fig. 2; compare Fig. 5a,d,g to Fig. 5b,e,h; Fig. S9). In comparison, the steepest clines were obtained in the NE scenario (Fig. 5c,f,i; Fig. S9), where the species had to adapt to novel environmental conditions for which no adapted genotypes were available within the range.

In the NoR model of phenotypic plasticity, the position of the perception trait (*g*_2_, sometimes also referred to as reference environment) had a strong influence on the spatial variation of the NoR slope (*g*_1_). The evolution of tolerance breadth was not affected by this parameter and evolved similar clines as the NoR model when *g*_2_ = 30. Otherwise, plasticity clines were reversed for lower values of *g*_2_, revealing the evolution of negative NoR slopes in RE and RS (Fig. 6a,b), but not in NE (Fig. 6c). This maladaptive plasticity did, however, not hinder adaptation to the local conditions and also allowed for phenotypic clines well aligned with the environmental gradient (Fig. 6d-e). It even favored colonization of new habitats in the NE scenario because moving the reference environment farther left on the range increased the phenotypic effects of allelic variation in plasticity (*g*_1_) (Fig. S10, Fig. A1).

**Figure 6:**
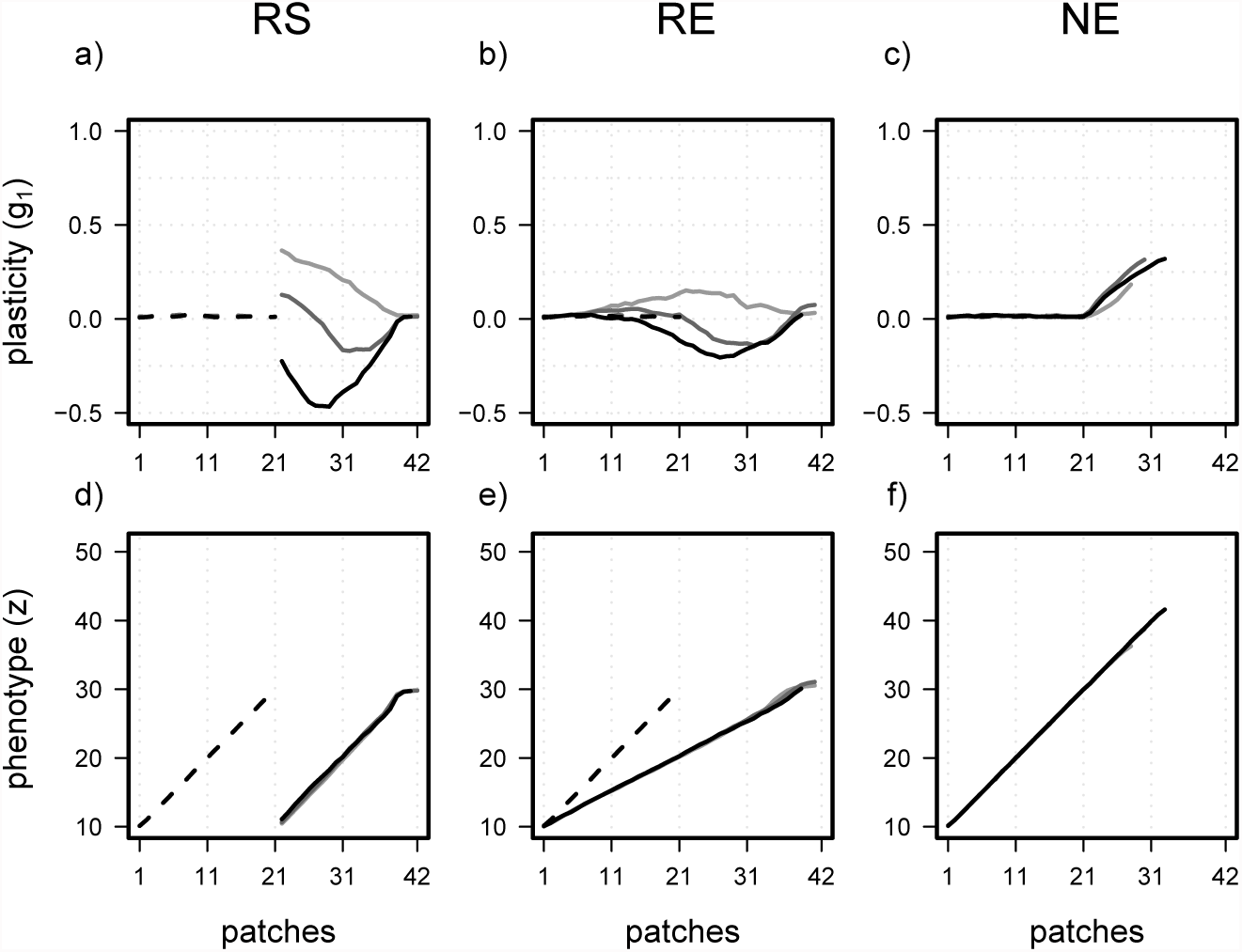
Phenotypic plasticity (*g*_1_, a-c) and the corresponding phenotypic values (*z*, d-f) are illustrated along the environmental gradient in dependence of the perception trait value (*g*_2_). Simulation results are given for high costs (*β* = 1.0), intermediate environmental variance (*σ*^2^(*e*) = 1), low migration rate (*m_c_* = 0.01), and for intermediate selection strength (*ω*^2^ = 4). Simulations were run with three distinct *g*_2_ values (*g*_2_ = 10 - black; *g*_2_ = 20 - dark gray; *g*_2_ = 30 - light gray).

## Discussion

Clines in environmental tolerance and phenotypic plasticity across a species’ range are widespread and considered a critical factor for the persistence of species facing environmental change (Valladares et al., 2014; Bennett et al., 2015). It has been suggested that tolerance clines evolve when species adapt to spatial clines of environmental variability, often resulting in higher tolerance at higher latitude (Addo-Bediako et al., 2000; Sunday et al., 2011), or when species modify their home range, during range expansion or range shift (Lancaster, 2016). We studied the evolution of tolerance and plasticity clines under the second less well covered hypothesis of species’ range evolution in three scenarios of range shift or expansion that were driven by environmental change or by niche evolution. We showed that in scenarios of range shift (RS) or range expansion (RE), without niche evolution, higher plasticity and tolerance evolved in parts of the range that experienced the longest history of environmental change, while lower plasticity was retained in areas reachable by pre-adapted genotypes. Therefore, we found the largest plasticity in trailing edge populations in RS, and in central populations in RE. In RS, trailing edge populations were occupied for the longest time and experienced the largest shift of their local conditions, while in RE, the central populations had the most favorable combination of extent and duration of past environmental change, provided migration was sufficiently limited. In contrast, in the scenario of niche evolution (NE) during invasion (or range expansion into new environments), the leading edge populations had the highest plasticity and tolerance because they were colonized by genotypes having to repeatedly adapt to novel environmental conditions. Migration favored plasticity and tolerance during colonization and thus reinforced the evolution of plasticity clines in the NE scenario, while it had an antagonistic effect on cline formation in the RS and RE scenarios because it favored the translocation of pre-adapted genotypes within the range, limiting the need for a plastic response. In sum, range dynamics can have a profound effect on the evolution of plasticity and tolerance and create clines that resemble empirical patterns. Our results thus confirm that range expansion driven by colonization of novel environments (NE) allows for the evolution of plasticity clines with a spatial increment similar to the pattern found for ectotherms’ latitudinal clines of thermal tolerance (Lancaster, 2016), while the other two evolutionary scenarios show results not usually reported in empirical studies. Of course, this does not discard the *climate variability hypothesis* but shows that an alternative explanation to clinal or latitudinal variation in plasticity and tolerance exists.

### Mechanisms behind the clines

During all of our range shift and range expansion simulations, plasticity and tolerance evolved in response to novel environmental conditions, experienced over time and across space. Plasticity clines then resulted when genotypes experienced different variability of local environments along the species’ range. However, in contrast to the *climate variability hypothesis*, the difference in variability was a consequence of the rate of environmental change in the different scenarios and of variation in migration rate instead of seasonal or between-year environmental fluctuations. Interestingly, range dynamics produced plasticity clines similar to those expected under climate variability only in the NE scenario where the environment was static but genotypes moved along the environmental gradient colonizing new habitats. In that case, plasticity becomes more advantageous at the colonization front where more plastic or tolerant individuals are selected. Costly plasticity may, however, strongly reduce the pace of colonization.

The evolved clines in environmental tolerance and phenotypic plasticity were transient and leveled out after the stabilization of the species’ distribution (Fig. S8), a process known as genetic assimilation (Waddington, 1953; Crispo, 2007; Lande, 2009). However, the decline in plasticity by genetic assimilation happens much slower than the build up of plasticity (Lande, 2009; Scheiner et al., 2017), such that the evolved clines outlasted the actual duration of the range dynamics by far. Our results therefore suggest long-lasting effects of species’ range dynamics on the tolerance and plasticity levels of species, despite their temporary nature.

The movement of genotypes, and thus the rate at which they are exposed to novel environmental conditions across space, was a key component of the evolution of plasticity clines in our simulations. Typically, plasticity was higher with increased migration rates in static environments, which also depended on the steepness of the fitness cline along the environmental gradient. Therefore, the equilibrium plasticity or tolerance level in static environments was a function of the spatial variation of fitness a migrating genotype was exposed to. Strong within-patch stabilizing selection (i.e., strong divergent selection) and high migration rates selected for high equilibrium levels of plasticity (see also Via and Lande, 1985; Scheiner, 1998; Sultan and Spencer, 2002). However, during range evolution, migration limited the formation of plasticity clines for two main reasons. First, high migration allowed for the translocation of previously locally adapted genotypes to the corresponding suitable habitat farther away on the shifted environmental gradient, reducing incentives to evolve higher plasticity or tolerance. This was particularly the case in the RS and RE scenarios, where plasticity clines were steeper for the lowest migration rates, and some plasticity evolved even at the front edge when genotypes could not catch up with their environment. Second, higher migration also imposed an evolutionary load on plasticity by bringing low-plasticity genotypes in environmentally variable populations, which resulted, for instance, in shallower plasticity clines at higher migration in the NE scenario (see Fig. 5f).

### Empirical patterns

Our finding of maximized tolerance and plasticity at the trailing edge in the RS scenario has, to the best of our knowledge, not been described empirically yet. Studies on within-species differences in plasticity or tolerance are rare in general (Valladares et al., 2014), and rear edge populations of dynamic species ranges (often at the warm margin) are understudied compared to those at the leading edge (Hampe and Petit, 2005; Thuiller et al., 2008). This rarity of empirical data is especially unfortunate given current ongoing climate changes. While a global temperature rise may progressively favor populations at the cold margin and allow them to expand polewards or upwards, tracking favorable conditions (Parmesan, 2006; Steinbauer et al., 2018), temperature increases are expected to negatively affect populations at the warm margins because they will experience novel environmental conditions outside of the species’ niche (Hampe and Petit, 2005; Kremer et al., 2012; Allendorf et al., 2013, p. 450-451). Thus, southern margin populations are supposed to be under stronger pressure to evolve or plastically respond to climate change (Duputié et al., 2015). Although we haven’t modeled niche expansion at the trailing edge, we expect evolving phenotypic plasticity to favor niche expansion also at the rear edge of species’ ranges and potentially rescue those populations from extinction.

The only study detecting patterns similar to those derived in our RE scenario is, as far as we know, Mägi et al. (2011) who found high morphological plasticity in the distribution center of *Agrimonia eupatoria*, while the closely related species *A. pilosa* at its distribution edge exhibited reduced plasticity for the same traits. However, Mägi et al. (2011) hypothesized that plasticity costs increased with environmental stress level causing lower plasticity in extreme habitats at the range edges. Our findings in RE scenarios add another potential explanation for these findings in the context of range expansion under environmental change and further empirical studies are necessary to discriminate between these alternative hypotheses. The scarcity of empirical evidence for high range center tolerance and plasticity is not surprising given the evolution of only shallow tolerance and plasticity clines in our simulations (Fig. 3, 4).

In contrast to the other two scenarios, our finding of elevated plasticity and tolerance at the leading edge in NE scenario is in line with several other theoretical (Roughgarden, 1972; Chevin and Lande, 2011; Lande, 2015) and empirical studies (Thomas et al., 2001; Matesanz et al., 2012; Lancaster, 2016), (but see Godoy et al., 2011; Palacio-López and Gianoli, 2011). Invasive species have been repeatedly found to have expanded their niche in novel habitats by evolving higher phenotypic plasticity and environmental tolerance (Molina-Montenegro and Naya, 2012; Atwater et al., 2017). Recently, Lancaster (2016) argued that this process could also explain latitudinal patterns of thermal tolerance in range-expanding ectotherms. Most of the range-expanding species in Lancaster (2016) were invasive species (16 out of 20), that rather expanded from low to high latitudes than vice versa. In line with this assumed expansion process, ectotherms were found to”overfill” their cold limit, i.e. were found beyond there previously measured cold margin, in a similar meta-analysis (Sunday et al., 2012). Interestingly, within-species increases in thermal tolerance and niche breadth with latitude were only observed at higher latitudes, but not (or only weaker) at lower latitudes (Lancaster, 2016; Papacostas and Freestone, 2016). In our simulations we found constant phenotypic plasticity and environmental tolerance in the part of the species’ range that served as a source for the colonization process, giving further support to the argument of Lancaster (2016).

### Comparing the tolerance curve and reaction norm approaches

We used two distinct approaches to simulate environmental tolerance evolution during species’ range dynamics–the tolerance curve and the reaction norm–and we observed that they do not always lead to the same qualitative results. Both approaches include two evolving quantitative traits that control the position of the environmental optimum (*t*_0_, *g*_0_) and the tolerance breadth (*t*_1_, *g*_1_). Deviations between the two approaches for the same parameter combination arose when the tolerance curve evolved broader tolerance breadths in response to environmental novelty, while the reaction norm approach resulted in maladaptive plasticity (negative *g*_1_) and thus smaller tolerance breadths (compare Fig. 4a with Fig. 6a). Conformity or nonconformity between tolerance curve and reaction norm evolution depended on the position of the perception trait value *g*_2_. The perception trait controls the phenotypic effects and the direction of plasticity (*g*_1_) evolution following the NoR equation (Equation 2), which illustrates that a phenotypic increase (*z*) is achieved by higher *g*_1_ values when *e* > *g*_2_, but by smaller or more negative *g*_1_ values when *e* < *g*_2_. To understand how maladaptive plasticity (negative *g*_1_) could be favored by evolution, it is necessary to distinguish between the fitness effect of plasticity in fluctuating environments and the contribution of plasticity to evolve novel phenotypes (see also Fig. A1c). In our simulations with a phenotypic optimum Θ increasing with the environment *e*, negative *g*_1_ values represented maladaptive plasticity because a single genotype that is adapted to a specific environment expresses phenotypes farther away from its new phenotypic optimum after an environmental perturbation. Negative slopes can, however, still be favored when they allow to express novel phenotypes during directional selection. Consequently, the evolution of maladaptive plasticity allowed to follow the phenotypic optimum over time but came at the expense of maladaptive responses of genotypes to local random environmental fluctuations.

We are aware that these consequences of the perception trait are entirely derived from geometrical reasoning, and a biological explanation of *g*_2_ is not immediately obvious. In fact, little attention has been paid to the evolutionary implications of the perception trait and many theoretical studies simply assumed *g*_2_ to be zero, to simplify the reaction norm equation. Studies on genetic *canalization* more explicitly refer to it as the *reference* environment where genetic and phenotypic variances are minimized (De Jong, 1990; Gavrilets and Scheiner, 1993; Lande, 2009; Chevin and Lande, 2011, see also Appendix 1). However, no model investigated the consequences of varying the reference environments on an environmental gradient, as we did here. One alternative would be to assume that locally adapted populations are canalized in their local environment and thus have each evolved a different value for the perception trait. Ergon and Ergon (2016) extended Lande’s model (Lande, 2009) for an *evolving* perception trait and showed that *g*_2_ could initially facilitate evolution towards novel phenotypic optima (when it evolves away from the novel *e*) and subsequently favor canalization (when *g*_2_ evolves towards the novel *e*). Ergon and Ergon (2016) justified their approach by arguing that the perception trait (*g*_2_) could be seen as a quantitative trait controlled by gene regulatory processes close to environmental cue perception, like cue activation thresholds for transduction elements. In contrast, the degree of plasticity (*g*_1_) might depend more on processes closer to phenotype expression that affect the sensitivity of gene regulation to changing transduction factors. Independently of its underlying genetic basis, empirical evidence of spatial heterogeneity and genetic variation in the perception trait are direly missing.

## Acknowledgement

We thank Lesley T. Lancaster, Thorbjørn H. Ergon, Luis-Miguel Chevin, Samuel M. Scheiner, Josh van Buskirk and Rassim Khelifa for valuable comments on the manuscript and lively discussions. Simulations were run on the UZH Science Cloud. MS and FG were funded by SNF grants PP00P3 144846 and PP00P3 176965.

## Author contributions

MS, RD, and FG designed the study. MS implemented evolving phenotypic plasticity and environmental tolerance in Nemo. MS and RD run the simulations and analyzed the data. FG and MS wrote the manuscript.

## Data accessibility

The Nemo source code, Nemo init files and summary of simulation results will be made available from the Dryad Digital Repository.

## Appendix

## App1 Potential consequences of the perception trait

The perception trait could have considerable effects on the evolvability of phenotypic plasticity as it controls the phenotypic effect sizes of *de novo* mutations and variance in *g*_1_. The perception trait could also lead to the evolution of maladaptive plasticity while adapting to novel environmental conditions. These effects become obvious from the reaction norm equation:
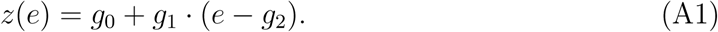

A mutation in the reaction norm slope has no phenotypic effect in the environment *e* = *g*_2_ when *g*_1_ · (*e* – *g*_2_) = 0. In other words, when only the NoR slope would evolve, there would be an invariant point where the reaction norm is fixed and is not affected by changes in *g*_1_ (Fig. A1). Instead, mutational effects of *g*_1_ increase with the distance between *e* and *g*_2_ (Fig. A1,b). Thus, the perception trait controls the environment dependence of the effect sizes of mutations as well as the additive genetic variance resulting from plasticity.

Based on the assumption that the perception trait itself could evolve, Ergon and Ergon (2016) showed that the perception trait *g*_2_ in response to environmental change first evolves away from the novel environmental value *e* (to allow for an elevated phenotypic variance and mutational effect sizes) and subsequently evolves towards the novel value *e* (to minimize the effects of (deleterious) mutations when the population reached the phenotypic optimum).

When the evolution of the perception trait is constrained also negative effects of *g*_2_ might result. Given that the perception is fixed to a value smaller than the average experienced environment (e.g., *g*_2_ = 10 with *e* = 20), a decrease in the environment with the associated decrease in the phenotypic optimum (when *e* = Θ) would favor the evolution of smaller or more negative slope values. While more negative slope values would allow to express phenotypes more closer to the novel optimum, they would lead to maladaptive plastic response in face of further environmental variation (Fig. A1c). We therefore argue that the adaptive value of plasticity could differ between the response to directional selection (adaptation to novel environments) and the effects during environmental fluctuations (e.g., temporal fluctuations).

**Figure A1:**
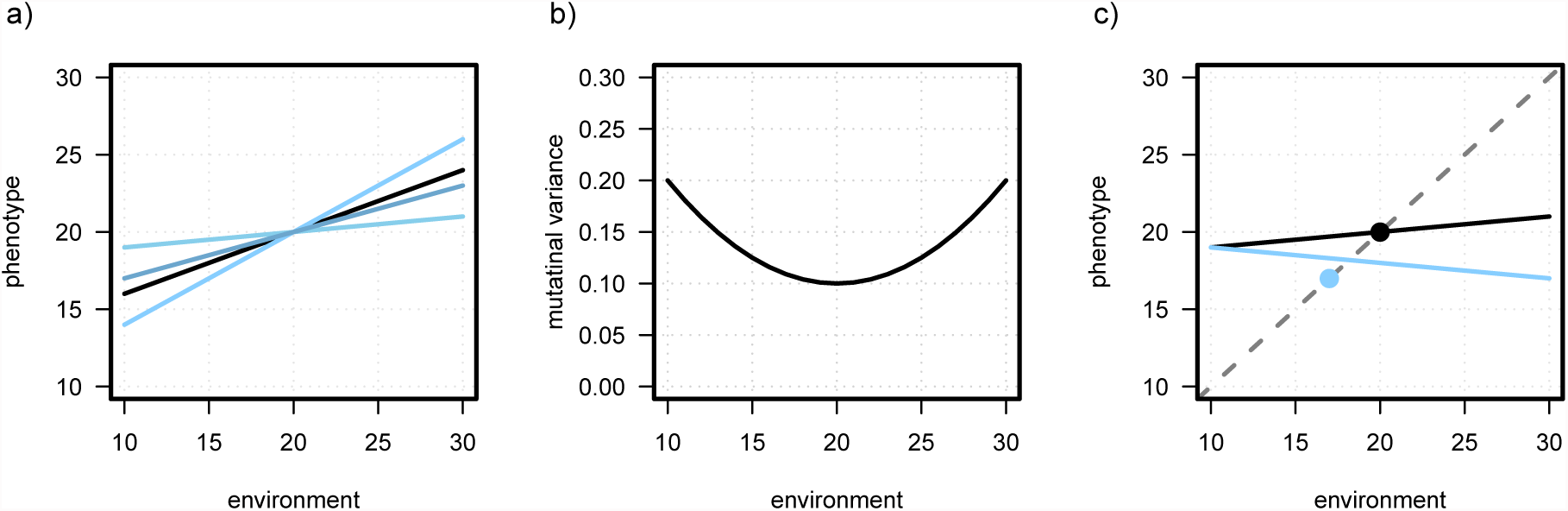
These graphs illustrate how the perception trait controls the phenotypic effects of *de novo* mutations in *g*_1_ (a,b), and how the perception trait could cause the evolution of maladaptive plasticity (c). Graph a) shows a genotypes’ reaction norm (black; *g*_0_ = 20, *g*_1_ = 0.4) and three potential mutant genotypes (blue) after a mutation in *g*_1_ (*g*_1_ = 0. 1, 0.3, 0.6) with a perception trait of *g*_2_ = 20. Graph b) visualizes the mutational variance resulting from mutations in the NoR intercept and NoR slope with a continuum-of-allele model. Mutational effects are picked from a bivariate normal distribution with variance of M[1,1]=0.1 for the intercept (*g*_0_) and M[2,2]=0.001 for plasticity (*g*_1_) with zero covariances (M[1,2]=M[2,1]=0). The results are shown for the analytical solution 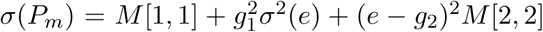 for the following wild type genotype: *g*_0_ = 20, *g*_1_ = 0, *g*_2_ = 20. Graph c) demonstrates how the perception trait could foster the evolution of maladaptive plasticity. The average reaction norm before environmental change (black line) allows to express a phenotype very close to the phenotypic optimum (black circle) in the environment *e* = 20. The dashed line represents the phenotypic optima (Θ) in dependence of the environment. Environmental change to *e* = 17 shifts the phenotypic optimum to 17 (blue circle) such that genotypes with lower (here negative) slope values are favored (e.g., blue line), when that the perception trait is fixed to *g*_2_ = 10,

## App2 How plasticity (*g*_1_) translates into tolerance (*t*_1_)

To translate the costs of phenotypic plasticity into costs of environmental tolerance, it is necessary to derive an equation of how *g*_1_ translates into *t*_1_. As a first step, we search for conditions when fitness depending on reaction norm parameters (*g*_0_, *g*_1_, *g*_2_) equals fitness depending on tolerance curve characteristics (*t*_0_, *t*_1_):
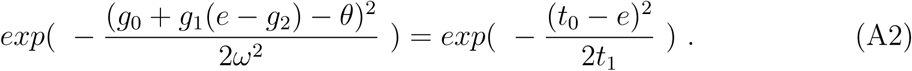

Multiplying this term by log() and −1, and simplifying the equation by assuming *t*_0_ = *g*_0_ = *g*_2_ = 0 as well as *e* = *θ* leads to:
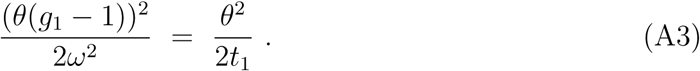

After dividing both sides by *θ*^2^ and multiplying by 2:
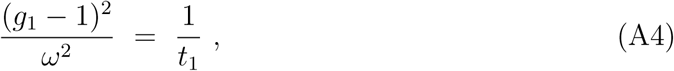

it is possible to solve for *t*_1_:
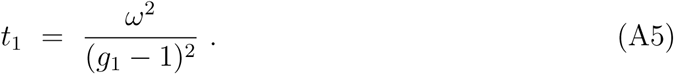

When solving for *g*_1_ following equation is derived (assuming 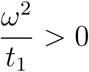):
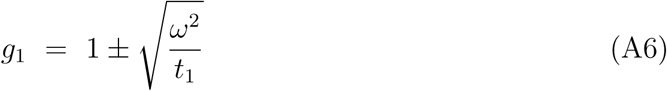

## Supplementary Material

**Table S1:** The following table shows the results of a linear model fitted to the levels of plasticity (*g*_1_, averaged over the species’ range) as dependent variable and the simulation parameters (*β* = 0. 0, 0. 5, 0.8,1.0,1.3, *m_c_* = 0, 0.001, 0.01, 0. 1, 0. 2, 0.4, *ω*^2^ = 1, 4,16) as predictive variables with a perception trait of *g*_2_=30 after burn-in simulations. While the sign and extend of the regression coefficients allow to deduce the parameters’ effects on average *g*_1_, p-values for simulation data are not meaningful and are not depicted here (White et al., 2014). The plasticity data were transformed with a logit function to obtain a more normally distributed response variable *g*_1_ (Warton and Hui, 2011). The adjusted-*R*^2^ was 0.86.

| Variable | Estimate | Std. Error |
| --- | --- | --- |
| (intercept) | 1.03 | 0.054 |
| costs of plasticity ($\beta$) | -0.80 | 0.047 |
| migration rate ($m_c$) | 0.23 | 0.153 |
| selection strength ($\omega^2$) | -0.02 | 0.003 |

**Figure S1:**
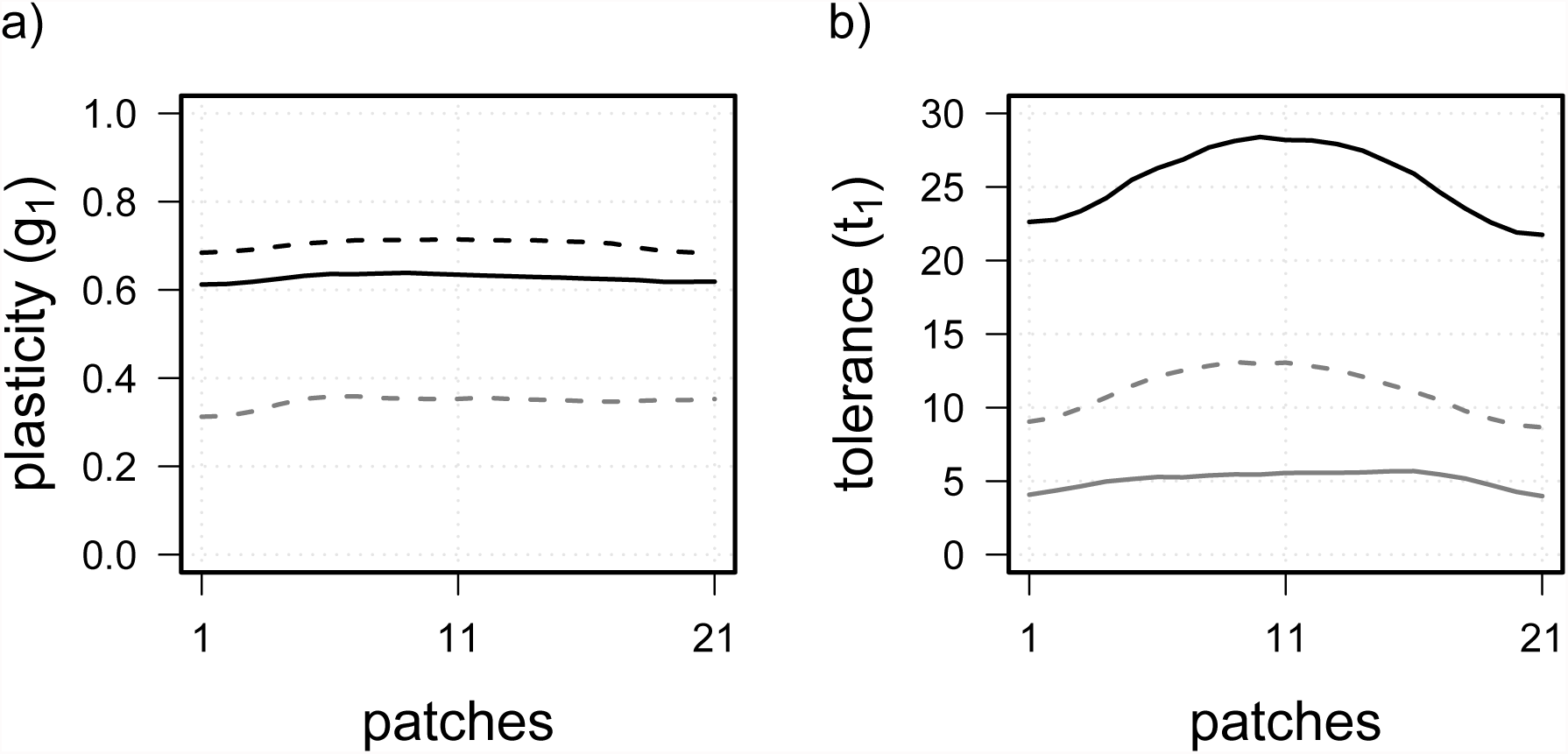
The two plots show the evolved levels of phenotypic plasticity (a) and environmental tolerance (b) after 100 000 generations (burn-in) with the most extreme edge effects observed in our simulations. Data derive from simulations with high migration rates (*m_c_* = 0.2 – *black*, *m_c_* = 0.4 – *gray*), strong selection (*ω*^2^ = 1), medium to high costs (*β* = 0.8 – *solid*, *β* = 1.0 – *dashed*), and *g*_2_ = 20.

**Figure S2:**
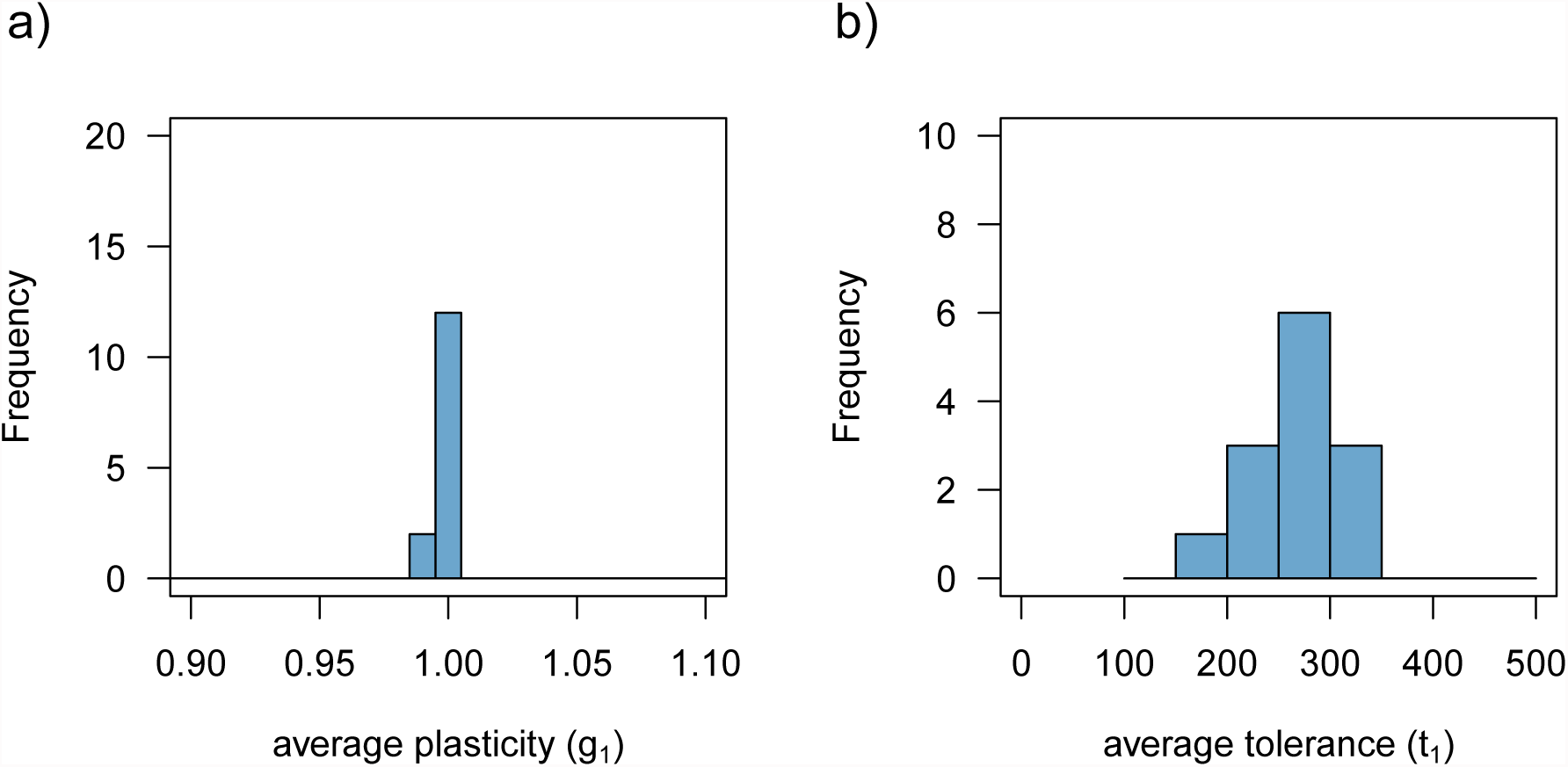
Histograms of simulation results for a) phenotypic plasticity (*g*_1_) and b) environmental tolerance (*t*_1_) for all three scenarios (RS, RE, NE) with zero costs (*β* = 0) after 100,000 generations in burn-in simulations. Plasticity and tolerance values are averaged over the occupied patches and replicates of the simulations for following parameter combinations: *ω*^2^ = 1, 4, 16, *m_c_* = 0.001, 0.01, 0.1, 0.2, 0.4, and *g*_2_ = 30. In absence of costs, perfect plasticity evolved in most scenarios except for those with low migration. Environmental tolerance evolved to values above 100 with no costs. From the 15 possible parameter combinations (*ω*^2^ × *m_c_*), repeated extinctions occurred and no data were obtained for one (for *g*_1_) and respectively two (for *t*_1_) parameter combinations with very low gene flow (*m_c_* = 0.001).

**Figure S3:**
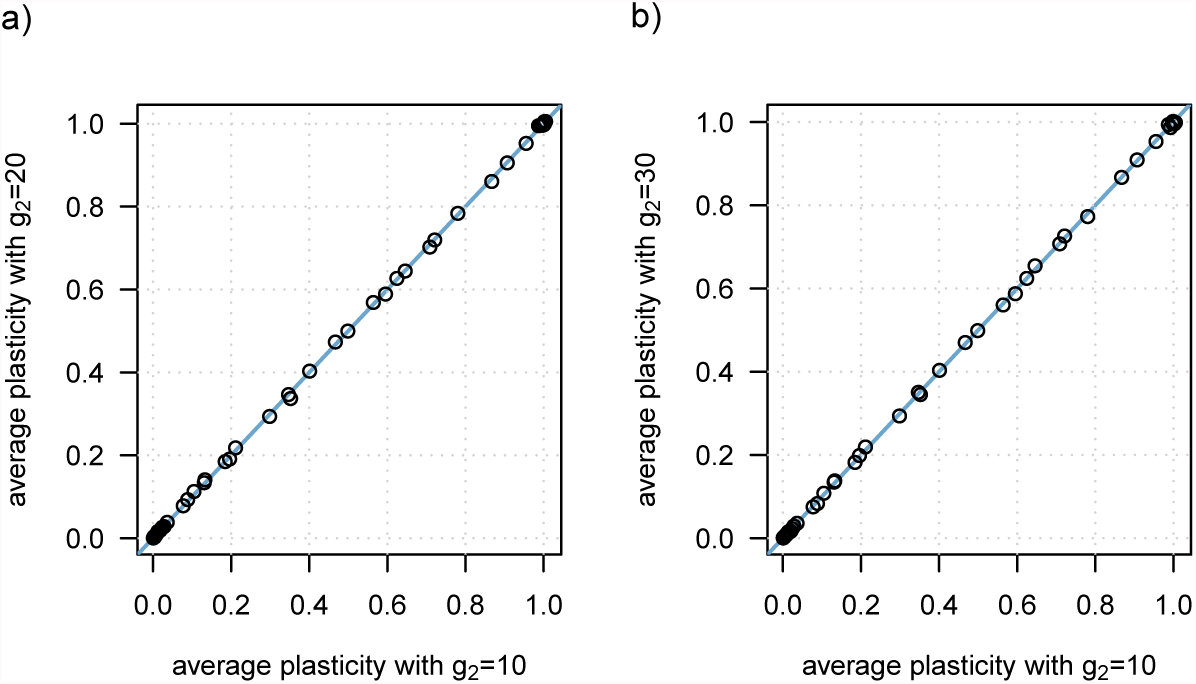
The two plots show the evolved levels of phenotypic plasticity (averaged over the species’ range and replicates) after 100,000 generations (burn-in) in dependence of the perception trait values. Results of parameter combinations with *g*_2_ = 10 are plotted against those with *g*_2_ = 20 in graph a) and against 92 = 30 in graph b). Data shown for the following parameter values: *ω*^2^ = 1, 4, 16, *β* = 0. 0, 0. 5, 0.8,1.0,1.3, and *m_c_* = 0.001, 0.01, 0. 1, 0. 2, 0.4. The 1:1 line is in blue.

**Figure S4:**
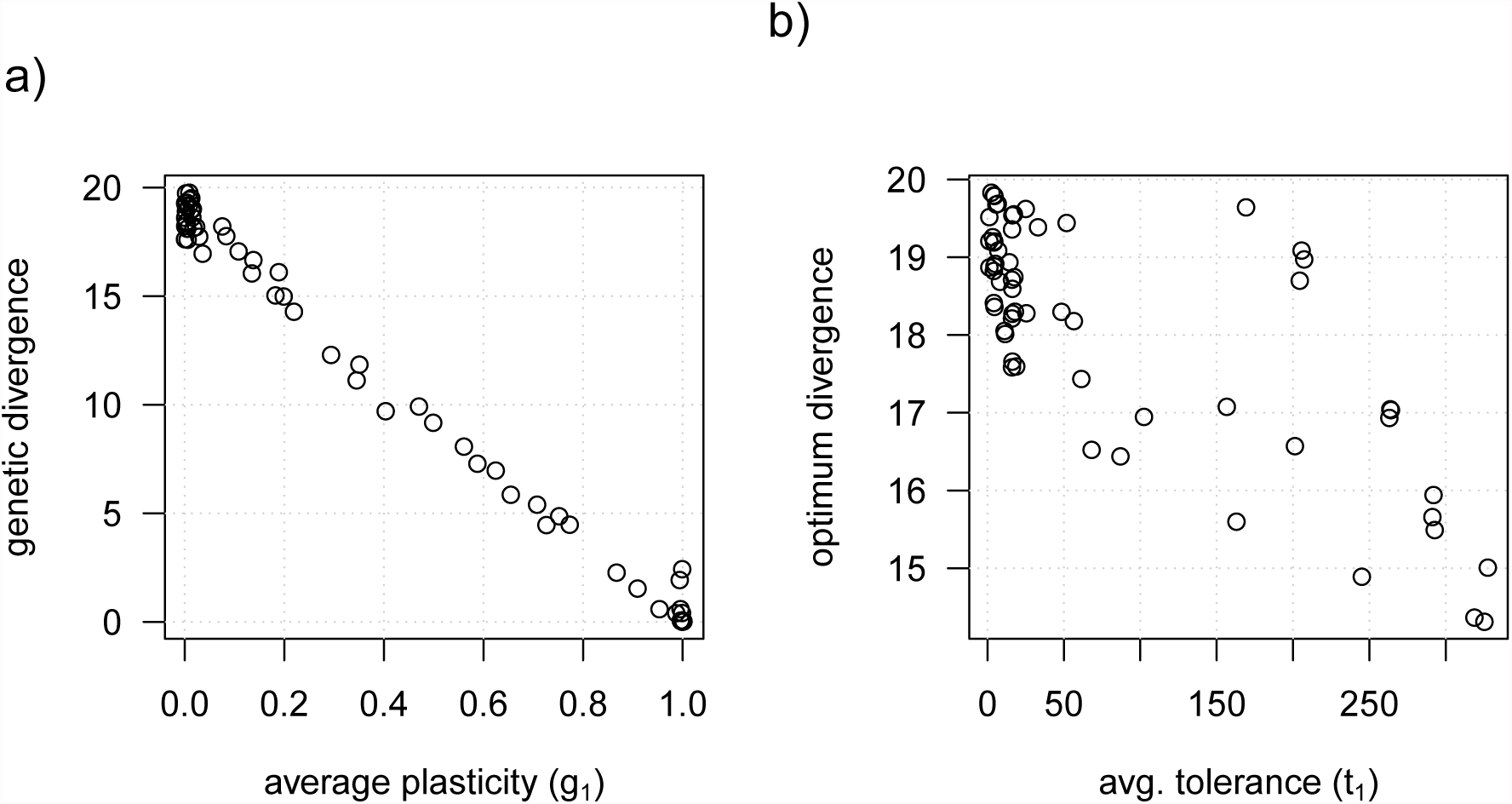
Graph a) shows that the levels of phenotypic plasticity (*g*_1_) after burn-in were strongly correlated with the genetic divergence along the range (*max*(*g*_0_) – *min*(*g*_0_)). Similarly, in graph b), higher environmental tolerance evolved together with smaller differences between the environmental optima along the species’ distribution (*max*(*t*_0_) – *min*(*t*_0_)). Results are plotted for following parameters: *ω*^2^ = 1, 4,16, *β* = 0.0, 0.5, 0.8,1.0,1.3, *m_c_* = 0.001, 0.01, 0.1, 0.2, 0.4, and *g*_2_ = 30.

**Figure S5:**
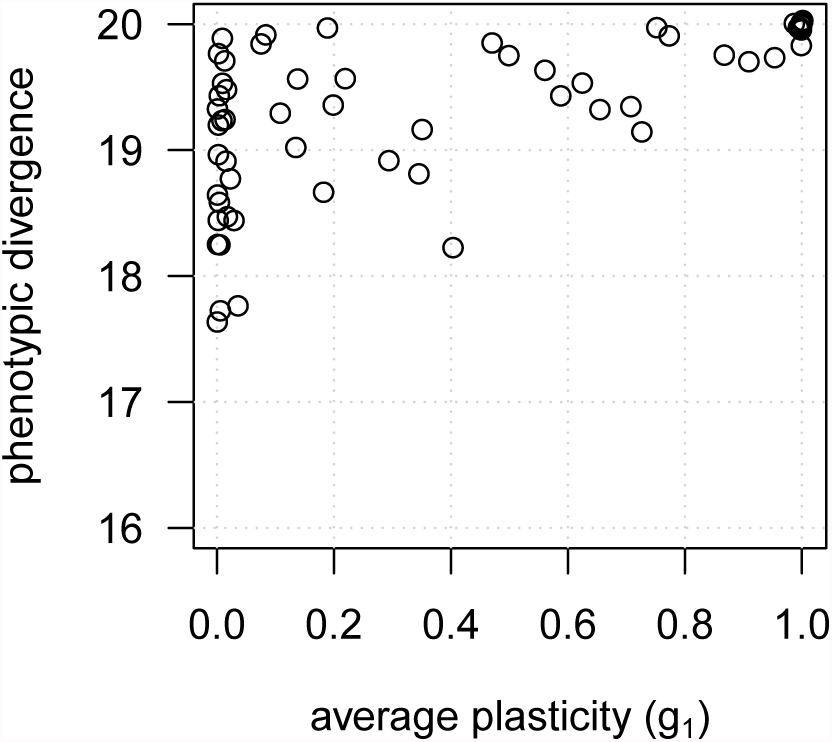
The graph shows the level of average phenotypic plasticity (*g*_1_) and the corresponding levels of phenotypic divergence (*max*(*z*) − *min*(*z*)) along the range after burn-in. High degrees of plasticity translated into large phenotypic differences along the range, with perfect plasticity allowing always for optimal phenotypic divergence. The plotted parameter combinations include *ω*^2^ = 1, 4, 16, *β* = 0.0, 0.5, 0.8, 1.0, 1.3, *m_c_* = 0.001, 0.01, 0.1, 0.2, 0.4, and *g*_2_ = 30.

**Figure S6:**
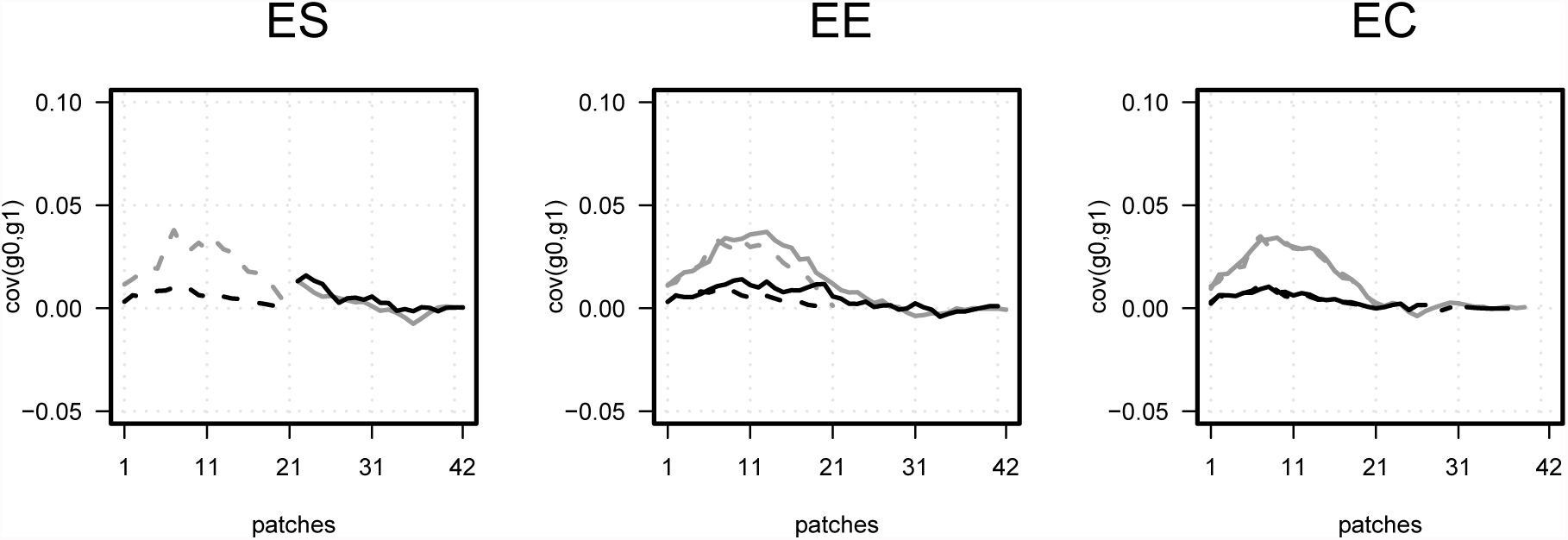
The covariance between individuals intercept and slope values within each patch before (dashed lines) and after (solid lines) environmental change (delta e = −0.1). For scenarios with *ω*^2^=4, *m_c_*=0.01, *g*_2_=30 for two different costs (*β*=0.5 - gray; *β*=1.0 - black)

**Figure S7:**
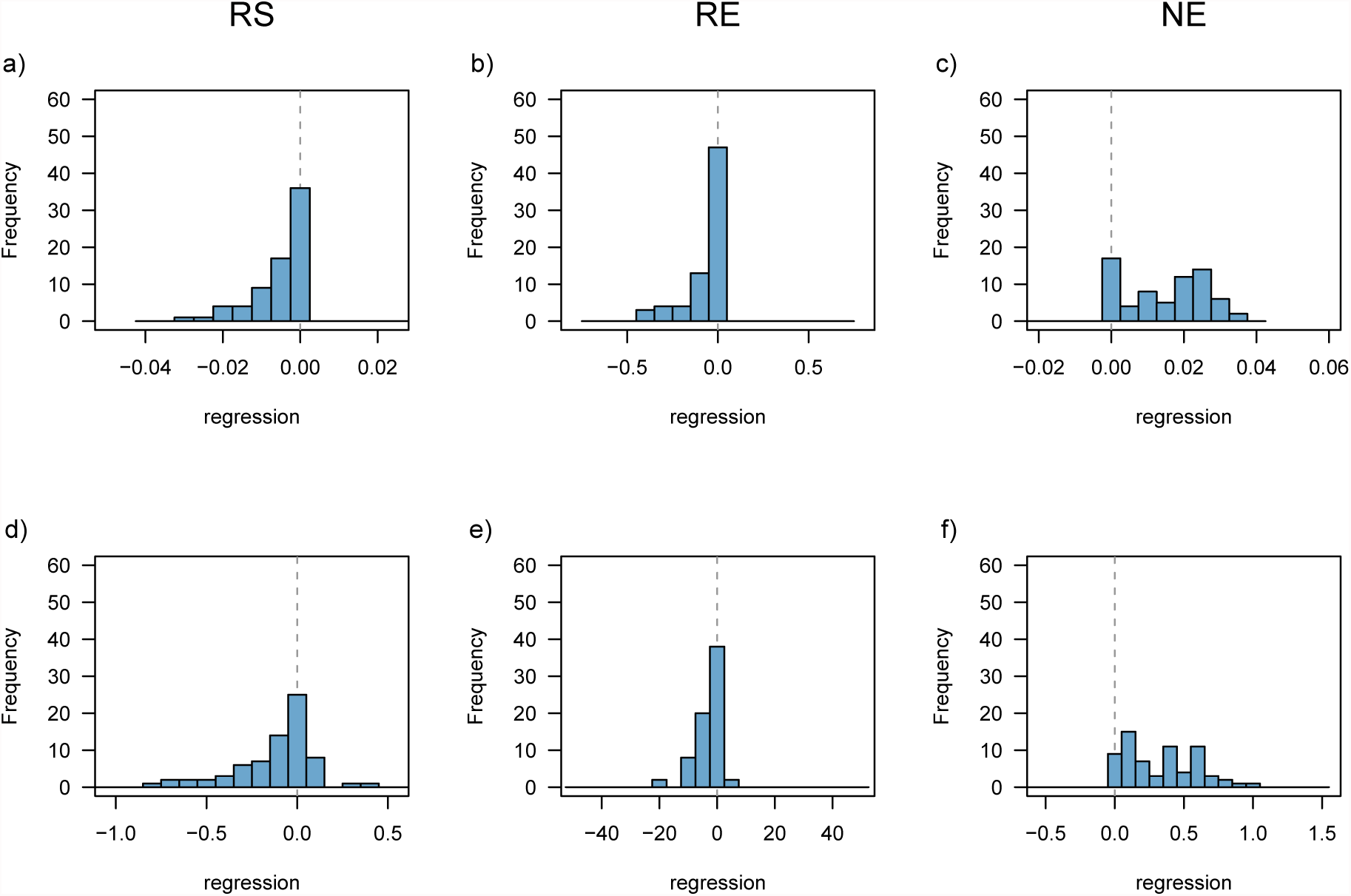
The histograms summarize the plasticity (a-c) and tolerance (d-f) clines evolved in our simulations after 210 generations of range shifts or range expansion. For RS scenarios, we performed a linear regression on the evolved *g*_1_ (a) and *t*_1_ (d) values along patches 22 to 42 and plotted the slope parameter of the linear model (second regression coefficient). Negative values indicate that plasticity decreased from the trailing edge towards the leading edge. We did the same for NE scenarios, when positive regression coefficients indicate an increase in plasticity towards the expansion front (c,f). For RE scenarios, we fitted a polynomial model along patches 1 to 42 for plasticity *g*_1_ (b) and tolerance *t*_1_ (e) and plotted the third regression coefficient with negative values indicating a humpback-shape fit. Regression coefficients were averaged over replicates for the following parameter combinations (*ω*^2^ = 1, 4, 16, *β* = 0.0, 0.5, 0.8,1.0,1.3, *m_c_* = 0.001, 0.01, 0.1, 0.2, 0.4, and *g*_2_ = 30), and were included in this plot when enough new patches have been colonized during the shift or expansion process.

**Figure S8:**
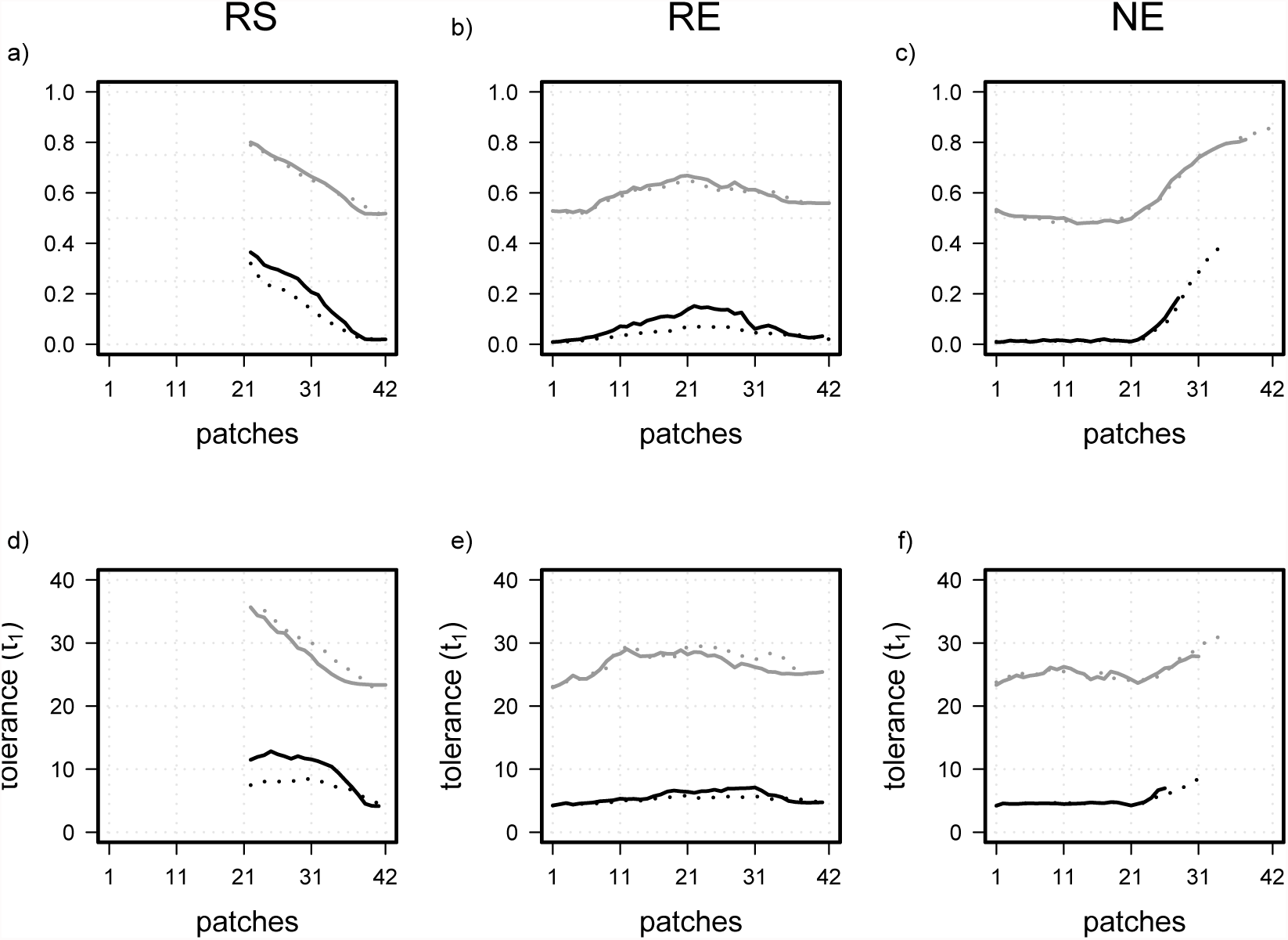
These figures illustrate the processes of genetic assimilation when phenotypic plasticity and environmental tolerance after 210 generations of range shift or range expansion (solid lines) levels out during additional 150 generations of constant environmental conditions (dotted lines). For scenarios with *ω*^2^ = 4, *m_c_* = 0.01, *g*_2_ = 30 for two different costs (*β* = 0.5 - gray; *β* = 1.0 - black). As the colonization process has not been finished during the first 210 generations in NE scenarios, there is no genetic assimilation happening (yet).

**Figure S9:**
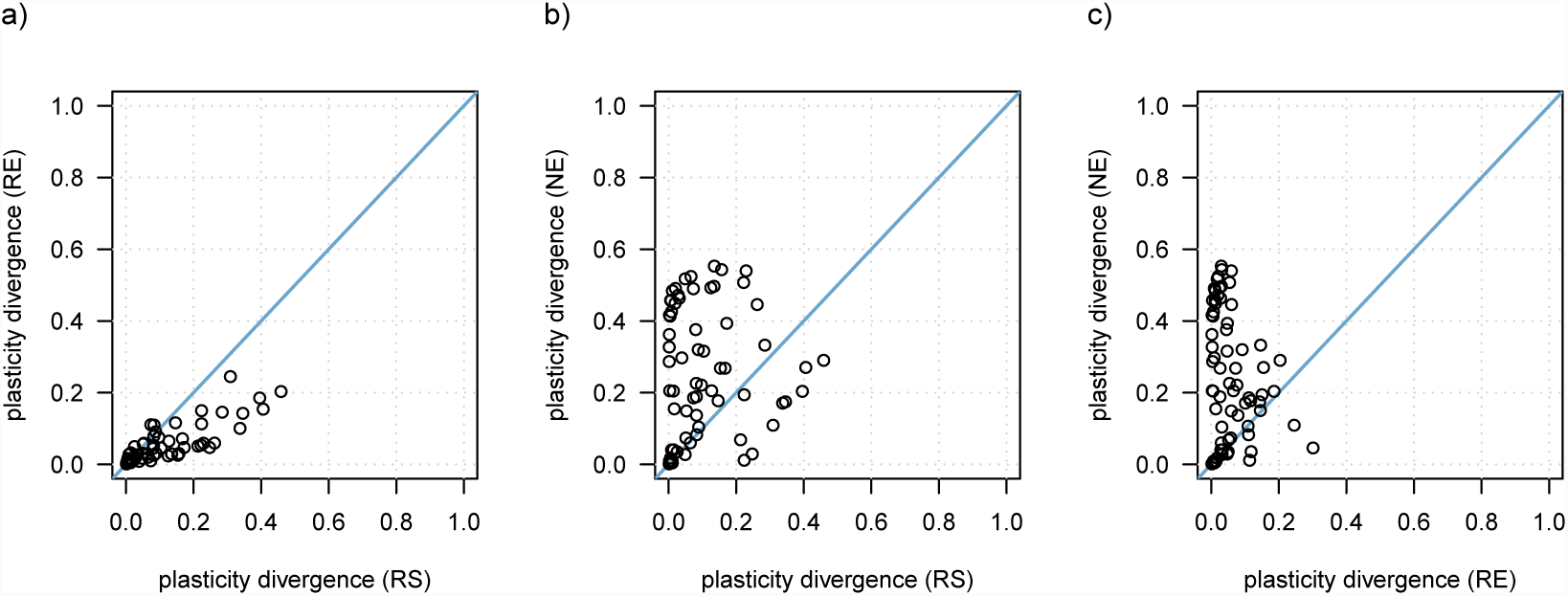
The divergence in plasticity (max(*g*_1_)-min(*g*_1_)) along the species’ range after range shift is compared between scenarios (RE, RS, NE) for all parameter combinations (*ω*^2^ = 1,4,16; *m_c_* = 0.001, 0.01, 0.1, 0.2, 0.4; *β* = 0,0, 0.5, 0.8, 1.0, 1.3) with a perception trait value of *g*_2_ = 30. Comparisons between scenarios are not included when at least in one of the two scenarios the species was not able to shift or expand its range.

**Figure S10:**
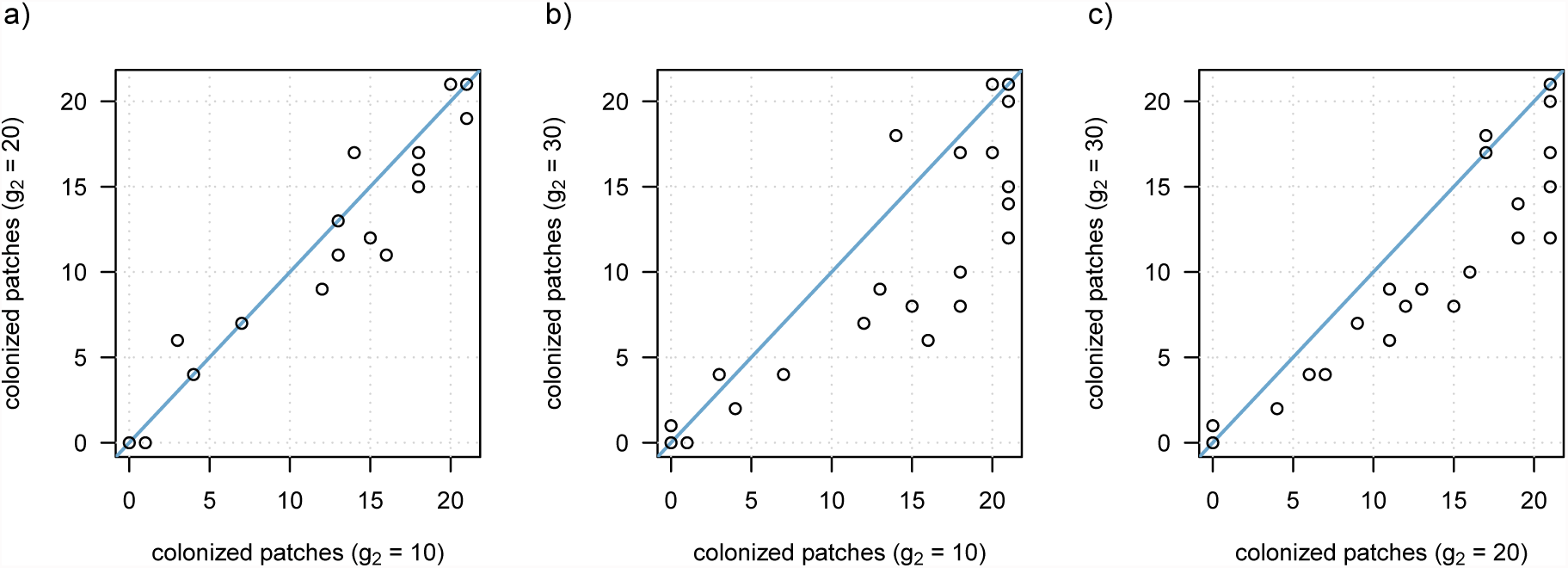
These figures illustrate the effect of the perception trait position on the ability to expand the distribution range and adapt to novel environmental conditions in NE scenarios. The number of colonized patches are plotted for all parameter combinations for the three combinations of perception trait values *g*_2_ = 10, *g*_2_ = 20, *g*_2_ = 30.

